# A human antibody that broadly neutralizes betacoronaviruses protects against SARS-CoV-2 by blocking the fusion machinery

**DOI:** 10.1101/2021.05.09.442808

**Authors:** Dora Pinto, Maximilian M. Sauer, Nadine Czudnochowski, Jun Siong Low, M. Alejandra Tortorici, Michael P. Housley, Julia Noack, Alexandra C. Walls, John E. Bowen, Barbara Guarino, Laura E. Rosen, Julia di Iulio, Josipa Jerak, Hannah Kaiser, Saiful Islam, Stefano Jaconi, Nicole Sprugasci, Katja Culap, Rana Abdelnabi, Caroline Foo, Lotte Coelmont, Istvan Bartha, Siro Bianchi, Chiara Silacci-Fregni, Jessica Bassi, Roberta Marzi, Eneida Vetti, Antonino Cassotta, Alessandro Ceschi, Paolo Ferrari, Pietro E. Cippà, Olivier Giannini, Samuele Ceruti, Christian Garzoni, Agostino Riva, Fabio Benigni, Elisabetta Cameroni, Luca Piccoli, Matteo S. Pizzuto, Megan Smithey, David Hong, Amalio Telenti, Florian A. Lempp, Johan Neyts, Colin Havenar-Daughton, Antonio Lanzavecchia, Federica Sallusto, Gyorgy Snell, Herbert W. Virgin, Martina Beltramello, Davide Corti, David Veesler

**Author notes:** These authors contributed equally (Dora Pinto, Maximilian M. Sauer, Nadine Czudnochowski, Jun Siong Low).

## Abstract

The repeated spillovers of β-coronaviruses in humans along with the rapid emergence of SARS-CoV-2 escape variants highlight the need to develop broad coronavirus therapeutics and vaccines. Five monoclonal antibodies (mAbs) were isolated from COVID-19 convalescent individuals and found to cross-react with multiple β-coronavirus spike (S) glycoproteins by targeting the stem helix. One of these mAbs, S2P6, cross-reacts with more than twenty human and animal β-coronavirus S glycoproteins and broadly neutralizes SARS-CoV-2 and pseudotyped viruses from the sarbecovirus, merbecovirus and embecovirus subgenera. Structural and functional studies delineate the molecular basis of S2P6 cross-reactivity and broad neutralization and indicate that this mAb blocks viral entry by inhibiting membrane fusion. S2P6 protects hamsters challenged with SARS-CoV-2 (including the B.1.351 variant of concern) through direct viral neutralization and Fc-mediated effector functions. Serological and B cell repertoire analyses indicate that antibodies targeting the stem helix are found in some convalescent donors and vaccinees but are predominantly of narrow specificity. Germline reversion of the identified cross-reactive mAbs revealed that their unmutated ancestors are specific for the endemic OC43 or HKU1 viruses and acquired enhanced affinity and breadth through somatic mutations. These data demonstrate that conserved epitopes in the coronavirus fusion machinery can be targeted by protective antibodies and provide a framework for structure-guided design of pan-β-coronavirus vaccines eliciting broad protection.

## Introduction

Severe acute respiratory syndrome coronavirus 2 (SARS-CoV-2), which is the causative agent of the coronavirus disease 2019 (COVID-19), emerged at the end of 2019 and seeded the ongoing pandemic (*1*). Coronaviruses are zoonotic pathogens found in avian and mammalian reservoirs, including bats, palm civets, racoon dogs, pangolins and pigs to cite a few. Repeated spillovers to humans demonstrates that these viruses have broad capability to spread between multiple phylogenetically distinct species. Over the past two decades, in addition to SARS-CoV-2, severe acute respiratory syndrome coronavirus (SARS-CoV) and Middle-East respiratory syndrome coronavirus (MERS-CoV) crossed the species barrier to humans leading to epidemics. All three highly pathogenic coronaviruses belong to the β-coronavirus genus: SARS-CoV-2 and SARS-CoV cluster within lineage B (*sarbecovirus* subgenus) and likely originated in bats whereas MERS-CoV belongs to lineage C (*merbecovirus* subgenus) and is transmitted to humans via contact with dromedary camels. Furthermore, four endemic coronaviruses cause common colds in humans: the HCoV-HKU1 and HCoV-OC43 β-coronaviruses in lineage A (*embecovirus* subgenus) and the more distantly related α-coronaviruses NL63 and 229E (*setracovirus* and *duvinacovirus* subgenus).

The coronavirus spike (S) glycoprotein promotes viral entry into host cells through an S_1_ subunit which engages host receptors and an S_2_ subunit involved in membrane fusion (*2*). The S_1_ subunit mediates receptor attachment, is the major target of (neutralizing) antibodies (Abs), and harbors the most sequence variability within the S glycoprotein (*3–6*). Accordingly, Abs binding to the receptor-binding domain (RBD) and N-terminal domain (NTD within the S_1_ subunit exert a selective pressure resulting in the emergence of new variants (*6–10*). The S_2_ subunit is more conserved than the S_1_ subunit among coronaviruses putatively due to the necessity to maintain the functionality of the fusion machinery and due to low immune pressure. The extensive glycan shield on the S_2_ subunit is also conserved among coronaviruses while retaining accessibility to functionally important regions, such as the fusion peptide (*11*). Abs binding to the S_2_ fusion machinery could potentially neutralize distantly related coronaviruses as reported for human and mouse mAbs (*12–14*) and observed for other viruses, such as influenza and HIV-1 (*15, 16*).

The emergence and rapid spread of SARS-CoV-2 has unequivocally demonstrated the need to develop interventions against coronaviruses, for this pandemic as well as for future spillover events. In the long term, the most effective drugs will be those with efficacy against a broad spectrum of coronaviruses, such as broadly neutralizing mAbs which target highly conserved S epitopes that are less likely to mutate and develop resistance. Furthermore, identifying conserved epitopes may guide the design of next generation pan β-coronavirus vaccines that could protect against both current and future emerging SARS-CoV-2 variants and other β-coronaviruses.

Here we describe the isolation of five mAbs that cross-react and broadly neutralize multiple human and animal β-coronaviruses by targeting the S glycoprotein stem helix. Structural and functional analyses of one of these mAbs (S2P6) indicate that inhibition of viral entry involves blocking membrane fusion which is the basis for broad β-coronavirus neutralization. S2P6 activates immune cell-dependent effector functions in vitro and protects hamsters challenged with SARS-CoV-2 Wuhan-1 and B.1.351 isolates leveraging both neutralization and effector functions. These findings pave the way for the development of pan β-coronavirus vaccines and therapeutics.

## Results

### Isolation of a broadly-neutralizing β-coronavirus mAb from a convalescent SARS-CoV-2 exposed individual

To identify mAbs targeting highly conserved regions of the S glycoprotein, we interrogated human IgG^+^ memory B cells from three COVID-19 convalescent donors. Five mAbs bound to the prefusion-stabilized S ectodomain trimers of viruses belonging to all three human-infecting β-coronavirus subgenera, i.e., *sarbecovirus* (SARS-CoV and SARS-CoV-2), *merbecovirus* (MERS-CoV) and *embecovirus* (OC43 and HKU1), but not to the human α-coronaviruses (229E and NL63) (**Fig. 1A-C** and **fig. S1A-B**). S2P6 and S2S43 mAbs were derived from two donors and use VH1-46*01 and D5-12*01 genes whereas the other 3 mAbs (P34D10, P34G12 and P34E3), derived from a third donor, are clonally related and use the VH3-30 gene (**fig. S1A**). Overall, the level of somatic mutations in these 5 mAbs in variable regions was moderate with nucleotide sequence identities to germline genes ranging from 86 to 97% (**fig. S1A**). The mAbs bound to both prefusion and post-fusion SARS-CoV-2 S with comparable apparent avidities, indicating that their cognate epitope is (at least partially) accessible in both conformational states of the S glycoprotein (**Fig. 1A** and **fig. S1B-C**).

**Fig. 1.**
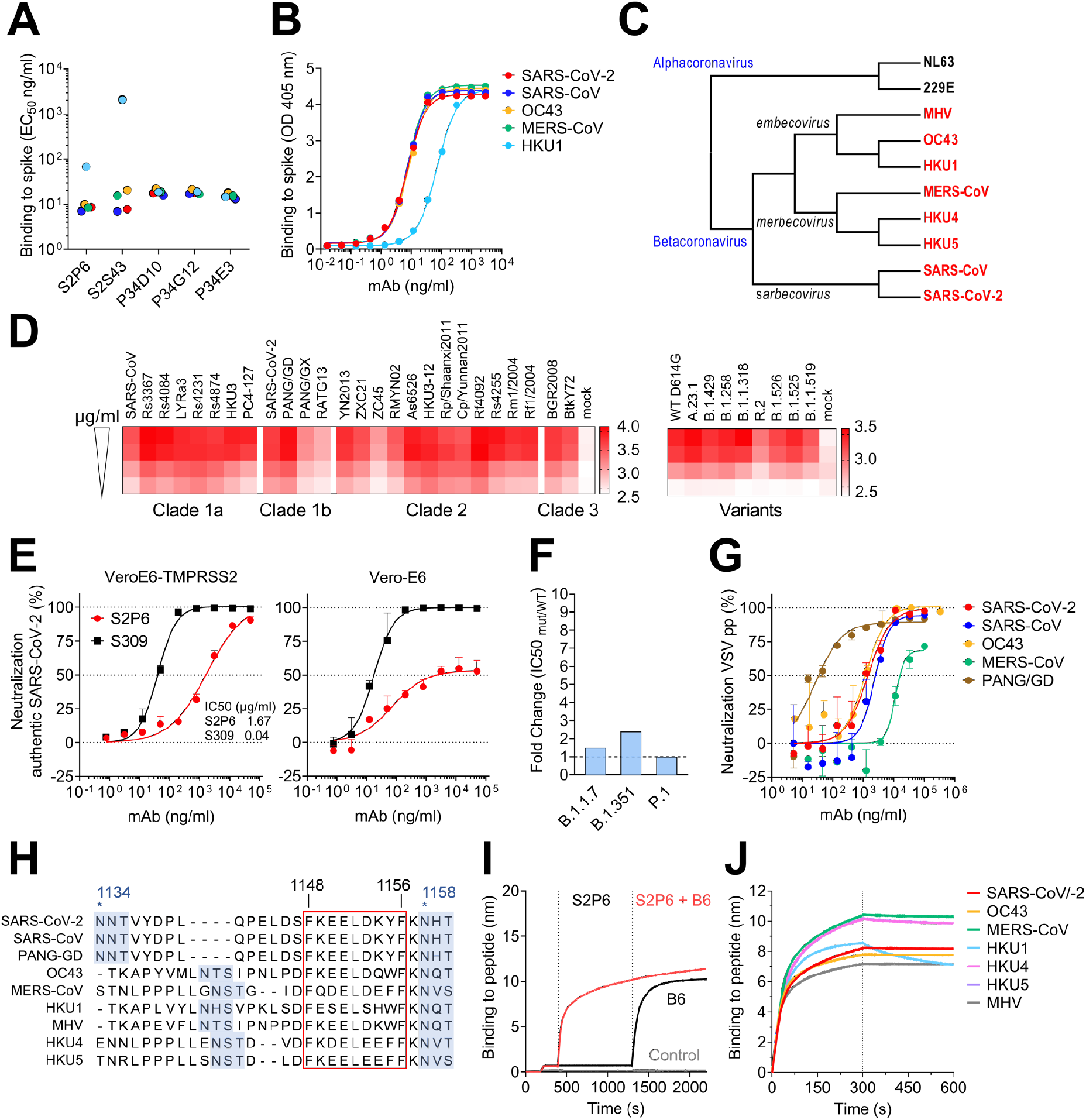
The S2P6 cross-reactive mAb broadly neutralizes β-coronaviruses from three subgenera. (**A-B**) Binding avidity (EC50) of 5 mAbs to prefusion coronavirus S trimer ectodomains as determined by ELISA (A). S2P6 binding curves from one representative experiment out of two is shown (B). (**C**) Cladogram of representative α- and β-coronavirus S glycoprotein amino acid sequences inferred via maximum likelihood analysis. β-coronaviruses are highlighted in red. (**D**) Flow cytometry analysis of S2P6 binding (from 10 to 0.22 μg/ml) to a panel of 26 S glycoproteins representative of all *sarbecovirus* clades (left) and 8 SARS-CoV-2 variants (right) displayed as a heat map of log MFI (mean fluorescent intensity). (**E**) Neutralization of authentic SARS-CoV-2 by S2P6 determined using VeroE6-TMPRSS2 (left) or Vero-E6 (right) cells. S309 mAb that binds RBD site IV (*21*) is included for comparison. Mean ± s.d. of triplicates from one representative experiment out of three is shown. (**F**) S2P6-mediated neutralization of SARS-CoV-2 B.1.1.7 S, B.1.351 S and P.1 S VSV pseudotypes represented as IC50 fold change relative to wildtype (D614G) S VSV pseudotype. (**G**) S2P6-mediated neutralization of VSV pseudotyped with various β-coronavirus S glycoproteins. Error bars indicate standard deviation of triplicates. IC50 values: 2.4 μg/ml, 1.4 μg/ml, 17.1 μg/ml, 1.3 μg/ml and 0.02 μg/ml for SARS-CoV, SARS-CoV-2, MERS-CoV, OC43 and PANG/GD, respectively. (**H**) Alignment of β-coronavirus stem helix region with the S2P6 epitope boxed. Residue numbering is shown according to SARS-CoV-2 S. N-linked glycosylation sequons are highlighted in blue. (**I**) Competition assay of S2P6/B6 binding to the identical stem helix peptide from SARS-CoV-2 and SARS-CoV (herein defined as SARS-CoV/-2). B6 binding in presence of S2P6 (red line), B6 binding in absence of S2P6 (black line) and the no SARS-CoV/-2 stem peptide control (grey line) is shown. (**J**) Kinetics of S2P6 binding to a panel of biotinylated β-coronavirus stem helix peptides immobilized at the surface of biolayer interferometry biosensors.

S2P6 was selected for further in-depth characterization and shown to bind to all full-length SARS-CoV-2 S variants tested and to 25 S glycoproteins (transiently expressed on the surface of ExpiCHO cells) representative of all *sarbecovirus* clades (**Fig. 1D** and **fig. S1D-E**). Using surface plasmon resonance (SPR), we found that S2P6 Fab fragment had the highest affinity for SARS-CoV-2 S and SARS-CoV S followed by MERS-CoV S and OC43 S with equilibrium dissociation constants (K_D_) of 7, 7, 12 and 16 nM, respectively (**fig. S1F-G**). S2P6 also bound to HKU1 S albeit with reduced affinity (K_D_ ~120 nM) (**fig. S1F**). The recognition of prefusion SARS-CoV-2 S by this mAb is pH dependent with higher binding affinity at pH 7, relative to pH 5, in both IgG and Fab formats (**fig. S1G**). Together, these data demonstrate the remarkable and efficient cross-reactivity of S2P6 towards all human-infecting β-coronaviruses.

To evaluate the neutralization potency and breadth of S2P6, we investigated its ability to inhibit entry of authentic SARS-CoV-2 into Vero-E6 cells in the presence or absence of the S-activating protease TMPRSS2. S2P6 completely neutralized infection of TMPRSS2-positive Vero-E6 cells but was less effective in neutralizing infection of Vero-E6 cells (**Fig. 1E**). Previous studies established that the main route of SARS-CoV-2 entry into cultured lung cells occurs through TMPRSS2-activated fusion with the cytoplasmic membrane (*17–19*). The more efficient S2P6-mediated neutralization of SARS-CoV-2 entry into Vero-E6 cells expressing TMPRSS2, relative to cells lacking this protease, would be consistent with reduced S2P6 binding at endosomal pH (**fig. S1G**) and suggests that S2P6 is maximally efficient towards the viral entry pathway associated with lung cell infection. We subsequently assessed S2P6-mediated neutralization of vesicular stomatitis virus (VSV) (*20*) pseudotyped with SARS-CoV-2 S of several variants of concern (VOC), including B.1.1.7, B.1.351 and P.1, and observed similar potency to that found against the parental SARS-CoV-2 D614G S (**Fig. 1F**). Moreover, S2P6 inhibited SARS-CoV S, Pangolin Guangdong 2019 (PANG-GD) S, MERS-CoV S and OC43 S VSV pseudotypes with IC50 values ranging from 0.02 to 17 μg/ml (**Fig. 1G**). S2P6 therefore features an unprecedented broad β-coronavirus neutralizing activity, including SARS-CoV-2 and SARS-CoV sarbecovirus clades as well as members of the merbecovirus and embecovirus subgenera.

To define the epitopes recognized by the identified mAbs, we performed peptide mapping experiments using 15-mer linear overlapping peptides (**fig. S2A**). All five mAbs bound to peptides containing the SARS-CoV-2 motif F_1148_KEELDKYF_1156_ (**Fig. 1H**) located in the stem helix within the S_2_ subunit. This region is strictly conserved in SARS-CoV, highly conserved among other β-coronaviruses, and overlaps with the epitopes of B6 (**Fig. 1I**) and 28D9, two mAbs which were previously identified following mouse immunization (*12, 13*). S2P6 bound efficiently to the stem helix peptides of the five β-coronaviruses that infect humans (albeit with a faster off-rate for HKU1) as well as of the MERS-CoV-related bat viruses (HKU4 and HKU5) and murine hepatitis virus (MHV) (**Fig. 1J** and **fig. S2B**). S2S43 exhibited similar overall binding to S2P6 with markedly weaker reactivity towards the HKU1, HKU4 and HKU5 peptides, whereas the three clonally related P34D10, P34G12, and P34E3 mAbs exhibited weaker or no binding to HKU4 and HKU5 peptides (**fig. S2B**).

### Structural basis for S2P6 binding to the conserved S glycoprotein stem helix

To unveil the molecular basis of the exceptional S2P6 neutralization breadth, we determined a cryo-EM structure of the SARS-CoV-2 S ectodomain trimer in complex with the Fab fragments of S2P6 and S2M11 (to lock the RBDs in the closed state (*22*)) at 4.1 Å overall resolution. The marked conformational dynamics of the region recognized by S2P6 limited the local resolution of the stem helix/Fab to approximately ~12 Å (**Fig. 2A**, **table S1** and **fig S3**). 3D classification of the cryo-EM data revealed incomplete Fab saturation (**fig. S3**). Our cryo-EM structure confirms that S2P6 recognizes the stem helix and suggests that the mAb disrupts its quaternary structure, which is presumed to form a 3-helix bundle in prefusion SARS-CoV-2 S (*3, 4, 12*). To overcome the limited resolution of the stem helix/Fab interface in the cryoEM structure, we determined a crystal structure of the S2P6 Fab in complex with the SARS-CoV-2 S stem helix peptide (residues 1146-1159) at 2.67 Å resolution (**Fig. 2B-D**, **fig. S4A** and **table S2**). The peptide folds as an amphipathic α-helix resolved for residues 1146 to 1159. S2P6 buries approximately 600 Å^2^ upon binding to its epitope using shape-complementarity and hydrogen bonding involving complementarity determining regions (CDR) H1-H3, L1 and L3. The light chain residues Y33, Y92, G93, P96, P97 and F99 as well as heavy chain residues Y33, I50, H57, T58, S59, P101, K102 and G103 form a deep groove in which the hydrophobic side of the stem helix docks via residues F1148, L1152, Y1155 and F1156. Binding specificity is provided through backbone hydrogen bonding of residues F1148_SARS-CoV-2_ and K1149_SARS-CoV-2_ with CDRH3 P101, side chain hydrogen bonding of residues E1151_SARS-CoV-2_ with CDRL1 Y33 and CDRL3 S94, D1153_SARS-CoV-2_ with CDRH1 Y33 side chains as well as Y1155_SARS-CoV-2 S_ with CDRH2 S59 through a water molecule (**Fig. 2C-D**). The contribution of each epitope residue was validated by single substitution scan analysis with most mutations at positions 1148, 1151-1153 and 1155-1156 abolishing S2P6 binding, highlighting the importance of these residues for binding (**fig. S5A**). A substitution scan analysis performed on the P34D10, P34G12 and P34E3 mAbs revealed a similar pattern of key interacting residues (**fig. S5A**). Residue Y1155 is conservatively substituted to F1238_MERS-CoV_ or W1237/1240_HKU1/OC43_ and residue D1153 is conserved in MERS-CoV and OC43 but mutated to S1235 for HKU1 (**fig. S5B**). The residue scan and structural results suggest that the reduced binding affinity of S2P6 for HKU1 S is (at least partially) due to the D1153_SARS-CoV-2_S1235_HKU1_ substitution which is expected to abrogate or dampen electrostatic interactions with the CDRH1 Y33 side chain hydroxyl (**Fig. 2C-D**).

**Fig. 2:**
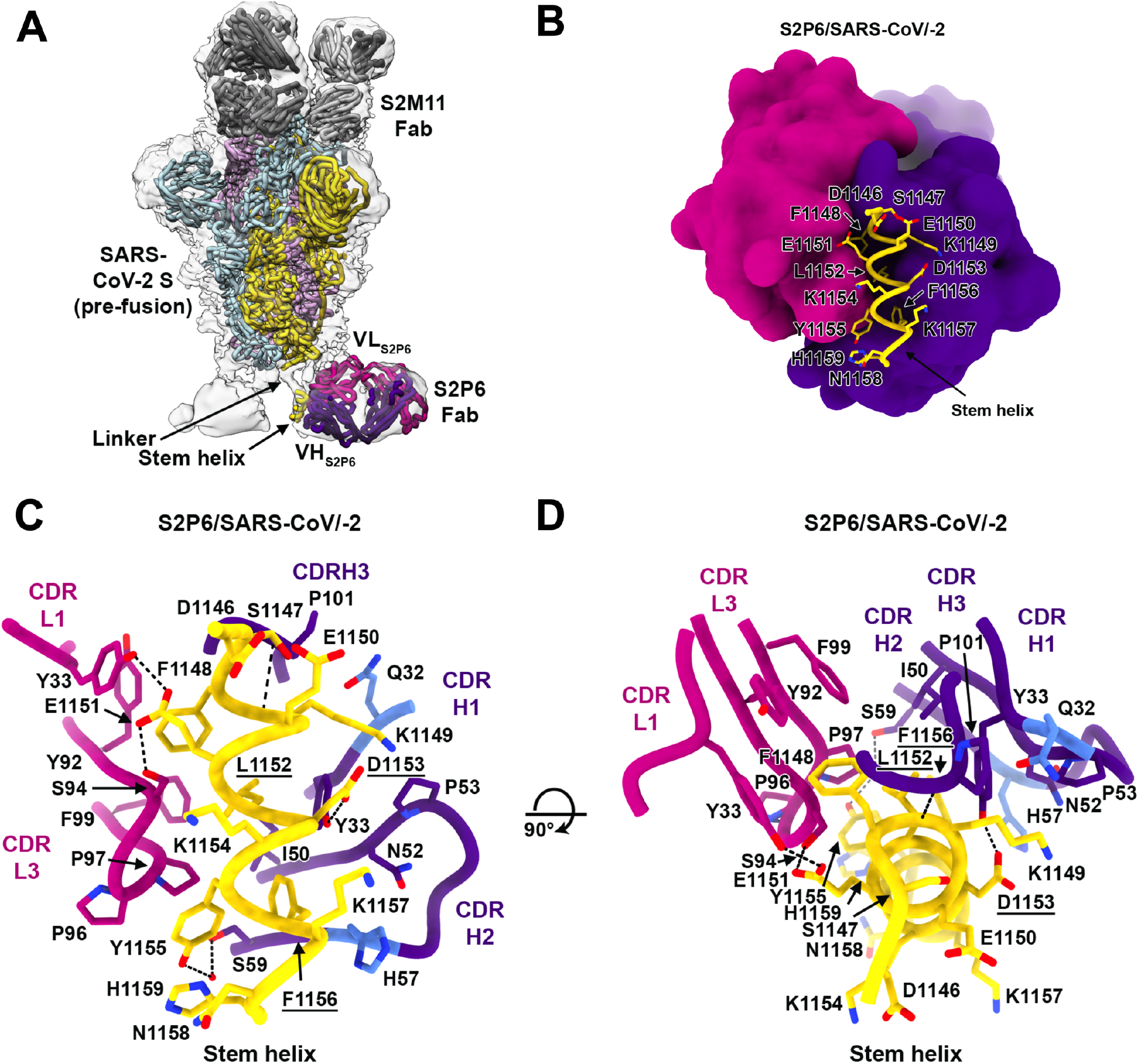
Structural basis for the broad S2P6 cross-reactivity with a conserved coronavirus stem helix peptide. (**A**) Composite model of the S2P6-bound SARS-CoV-2 S cryoEM structure and of the S2P6-bound stem helix peptide crystal structure docked in the cryoEM map (transparent gray surface). SARS-CoV-2 S protomers are colored pink, cyan and gold, the S2P6 Fab heavy and light chains are colored purple and magenta and the S2M11 Fab heavy and light chains are colored dark and light gray, respectively. (**B**) Ribbon diagram of the S2P6 Fab (surface rendering) in complex with the SARS-CoV-2 S stem helix peptide (yellow ribbon with side chains rendered as sticks and labeled). (**C-D**) Ribbon diagram in two orthogonal orientations of the S2P6 Fab bound to the SARS-CoV-2 S stem helix peptide showing a conserved network of interactions. Only key interface residues and the S2P6 CDR loops are shown for clarity. Residues Q32 and H57, that are mutated during affinity maturation of the S2P6 heavy chain, are colored blue. Hydrogen bonds are indicated with dashed lines. Residues substituted in the escape mutants isolated are underlined.

Although S2P6 and B6 recognize a similar epitope (**fig. S4B-C**), they bind with opposite orientations of the heavy and light chains relative to the stem helix and the S2P6-bound structure resolves 3 additional C-terminal peptide residues (1156-1159) compared to the B6-bound structure (1147-1156) (**fig. S4B-C**). Superposition of both structures based on the stem helix reveals that B6 CDRH2 would sterically clash with H1159_SARS-CoV-2_, putatively explaining the broader cross-reactivity of S2P6 over B6 (**fig. S4B-C**).

To further validate our structural data, we carried out viral escape mutant selection in vitro in the presence of S2P6 using a replication-competent VSV-SARS-CoV-2 S chimeric virus (*23*). After two passages, virus neutralization by S2P6 was abrogated and deep sequencing revealed the emergence of five distinct resistance mutations: L1152F, D1153N/G/A and F1156L, which are consistent with the structural data and substitution scan analysis (**fig. S5A**). These mutations have been detected with very low frequency in circulating SARS-CoV-2 isolates (146 out of 1,217,814 sequences as of April 30, 2021). The isolation of escape mutants in a VSV-SARS-CoV-2 S chimeric virus system contrasts with the high conservation of the stem helix and suggest that a selection pressure not recapitulated here could limit accumulation of mutations in this epitope in the authentic virus.

### Stem helix-targeting mAbs inhibit S-mediated membrane fusion

The S stem helix forms a 3-helix bundle in many prefusion cryoEM structures of SARS-CoV-2 S and SARS-CoV S (*3, 4, 11, 24–26*). In contrast, the S2P6/S2M11/SARS-CoV-2 S cryoEM structure suggests that the quaternary organization of the stem is disrupted (**Fig. 2A**) as supported by the observation that S2P6 binds to the hydrophobic face of the stem helix which is expected to be mostly buried through homo-oligomeric interactions in prefusion S, and that may be only transiently available for Ab binding. Although we observed S2P6 binding to post-fusion SARS-CoV-2 S (**fig. S1C**), the epitope recognized is buried at the interface with the other two protomers of the rod-shaped trimer and is therefore not expected to be fully accessible (**fig. S4D**) (*12, 27–29*). Based on these data, we hypothesized that S2P6 binding to S sterically interferes with the conformational changes leading to membrane fusion, as observed for B6 (*12*) and 27D9 (*13*).

To validate the inferred mechanism of S2P6-mediated broad coronavirus neutralization, we first showed that S2P6 binding did not block engagement of SARS-CoV-2 S by ACE2 using ELISA, as expected based on the remote location of its epitope from the RBD (**Fig. 3A**). S2P6, however, blocked cell-cell fusion between Vero-E6 cells transfected with full-length SARS-CoV-2 S as effectively as the S2M11 mAb which locks SARS-CoV-2 S in the closed state (*22*) (**Fig. 3B**). We previously described mAbs targeting RBD antigenic site Ia (e.g. S2E12) and IIa (e.g. S2X259 or S2X35) which can mimic receptor attachment and prematurely trigger fusogenic S conformational changes (*11, 30, 31*). Accordingly, S2P6 at concentrations as low as 1 ng/ml abrogated the formation of syncytia mediated by S2E12 (*30*). Collectively, these results suggest that the main mechanism of S2P6 neutralization is to prevent viral entry via inhibition of membrane fusion resulting from impeding S fusogenic rearrangements (**Fig. 3C**).

**Fig. 3.**
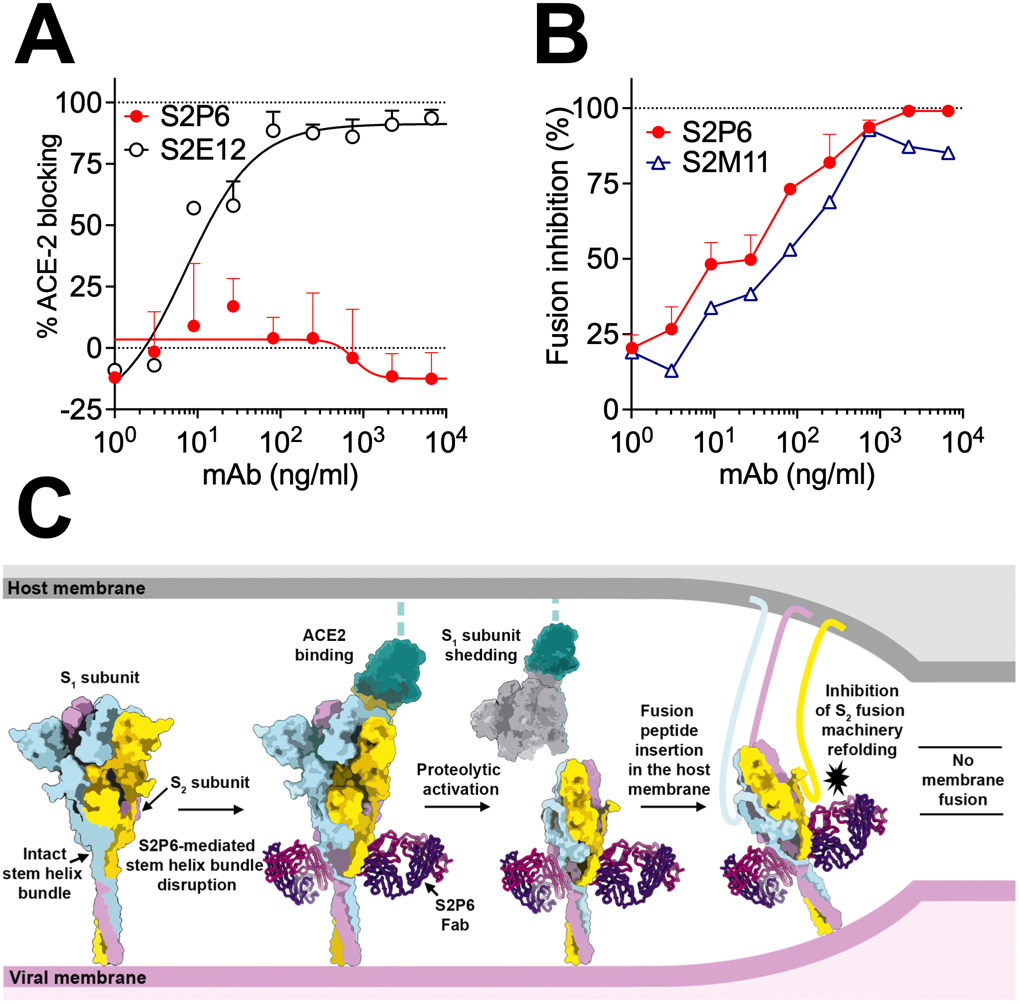
S2P6 binding disrupts the stem helix bundle and sterically inhibits membrane fusion. (**A**) SARS-CoV-2 S binding to ACE2 in the presence of mAb S2P6 analyzed by ELISA. S2E12 was included as a positive control. (**B**) S2P6 inhibition of cell-cell fusion using Vero-E6 cells transfected with SARS-CoV-2 S. S2M11 was included as a positive control. Inhibition of fusion values are normalized to the percentage of fusion without mAb (100%) and to that of fusion of non-transfected cells (0%). (**C**) Proposed mechanism of inhibition mediated by the S2P6 mAb. S2P6 binds to the hydrophobic core of the stem helix bundle and disrupts its quaternary structure. S2P6 binding likely prevents S_2_ subunit refolding from the pre-to the post-fusion state and blocks viral entry.

### S2P6-mediated protection is enhanced by Fc-mediated effector functions in hamsters

Fc-mediated effector functions can contribute to in vivo protection by promoting viral clearance and anti-viral immune responses (*32–35*). We analyzed the ability of S2P6 to trigger activation of FcγRIIa and FcγRIIIa, as well as to exert Fc effector functions in vitro. S2P6 promoted moderate dose-dependent FcγRIIa and FcγRIIIa mediated signaling using a luciferase reporter assay (**Fig. 4A**). S2P6 promoted robust activation of Ab-dependent cell cytotoxicity (ADCC), to levels comparable to those observed with the S309 mAb (*21*), following incubation of SARS-CoV-2 S expressing CHO-K1 target cells with human peripheral blood mononuclear cells (PBMCs) (**Fig. 4B**). S2P6 also triggered Ab-dependent cellular phagocytosis (ADCP) activity using CellTrace-Violet-labelled PBMCs as phagocytic cells, and SARS-CoV-2 S expressing CHO (CHO-S) as target cells (**Fig. 4C**). Finally, S2P6 did not promote complement-dependent cytotoxicity (CDC) (**Fig. 4D**) indicating that S2P6 Fc-mediated effector functions, but not complement activation, might participate in viral control in vivo.

**Fig. 4.**
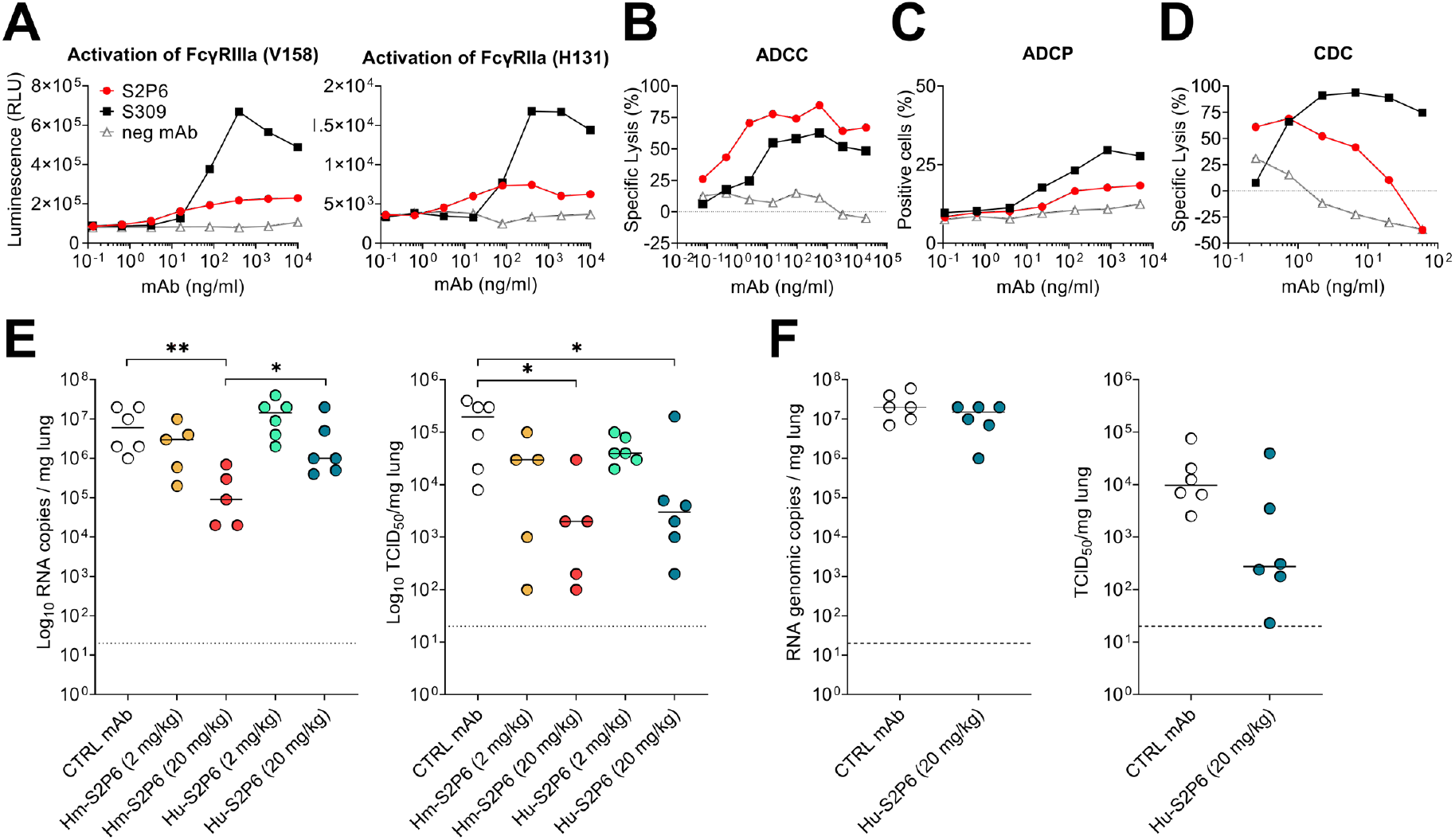
S2P6 activates effector functions and protects Syrian hamsters from SARS-CoV-2 challenge. (**A**) NFAT-driven luciferase signal induced in Jurkat cells stably expressing FcyRIIIa (V158, left) or FcγRIIa (V131, right) upon S2P6 binding to full-length wild-type SARS-CoV-2 S expressed at the surface of CHO target cells. S309 is included as positive control. RLU, relative luminescence unit. (**B**) mAb-mediated ADCC using SARS-CoV-2 CHO-K1 cells (genetically engineered to stably express a HaloTag-HiBit-tagged) as target cells and PBMC as effector cells. The magnitude of NK cell-mediated killing is expressed as percentage of specific lysis. (**C**) mAb-mediated ADCP using Cell-Trace-Violet-labelled PBMCs as a source of phagocytic cells (monocytes) and PKH67-fluorescently labeled S-expressing CHO cells as target cells. The *y*-axis indicates percentage of monocytes double-positive for anti-CD14 (monocyte) marker and PKH67. (**D**) Lysis of SARS-CoV-2 S stably transfected CHO cells by mAbs in the presence of complement (CDC assay). (**E**) Syrian hamsters were administered with the indicated amount of S2P6 mAb harboring either a hamster (Hm-S2P6) or a human (Hu-S2P6) constant region before intranasal challenge with prototypic SARS-CoV-2 (Wuhan-1 related). An irrelevant mAb (MGH2 against CSP of *P. falciparum*) at 20 mg/kg was used as negative control (*37*). Shown are viral RNA loads (left) and replicating virus titers (right). (**F**) Prophylactic administration of 20 mg/kg of human S2P6 in hamsters challenged with SARS-CoV-2 B.1.351 VOC. Shown are viral RNA loads (left) and replicating virus titers (right). * P<0.05, ** P<0.01 Mann-Whitney test.

We next evaluated the prophylactic activity of S2P6 against challenge with the prototypic (Wuhan-1 related) SARS-CoV-2 in a Syrian hamster model (*36*). As we previously showed that human IgG1s poorly recognize FcγRs (*30*), we compared S2P6 harboring a human IgG1 (Hu-S2P6) or a hamster IgG2a constant region (Hm-S2P6), the latter enabling optimal interactions with hamster FcγRs. Two different doses of Hu-S2P6 or Hm-S2P6 were administered 24 hours prior to intranasal SARS-CoV-2 challenge and the lungs of the animals were assessed 4 days post-infection for viral RNA load and replicating virus. Hm-S2P6 administered at 20 mg/kg reduced viral RNA copies and replicating viral titers in the lungs of animals by two orders of magnitude relative to a control mAb (**Fig. 4E**). Moreover, Hm-S2P6 at 20 mg/kg reduced viral RNA copies detected in the lungs to levels significantly lower than those observed with Hu-S2P6, suggesting a beneficial effect of S2P6 effector functions in vivo. Based on the comparable S2P6 neutralization potencies towards SARS-CoV-2 VOC observed in vitro (**Fig. 1F**), we set out to assess the protective efficacy of S2P6 in hamsters challenged with SARS-CoV-2 B.1.351. Prophylactic administration of Hu-S2P6 at 20 mg/kg reduced replicating viral titers in the lungs (but not RNA copy numbers) by ~1.5 order of magnitude relative to the control group, in line with the strict conservation of the stem helix epitope in all VOC identified to date (**Fig 4F**).

Collectively, these findings demonstrate that Abs targeting a highly conserved epitope in the S fusion machinery can trigger Fc-mediated ADCC and ADCP in vitro and protect against SARS-CoV-2 challenge by leveraging both neutralization and effector functions in vivo.

### Natural infection or vaccination predominantly elicit stem helix-directed Abs of narrow specificities

To understand how frequently stem helix-specific Abs are elicited, we performed serological analysis using plasma samples from pre-pandemic, COVID-19 convalescent, and COVID-19 vaccinated individuals (**Fig. 5A** and **fig. S6**). We determined the titers of plasma IgG binding to stem helix peptides of SARS-CoV-2/SARS-CoV (SARS-CoV/-2), OC43, MERS-CoV, HKU1, HKU4 and HKU5. We did not observe plasma Abs binding to stem helix peptides in prepandemic samples, except for HKU1, probably reflecting prior infection with this virus in this cohort. Conversely, stem helix-specific Abs were found at low frequency in individuals previously infected with SARS-CoV-2 or receiving two doses of mRNA vaccines (**Fig. 5A**). Overall, these data show that plasma Ab responses to the stem helix are elicited upon SARS-CoV-2 infection or vaccination although they are relatively rare.

**Fig. 5.**
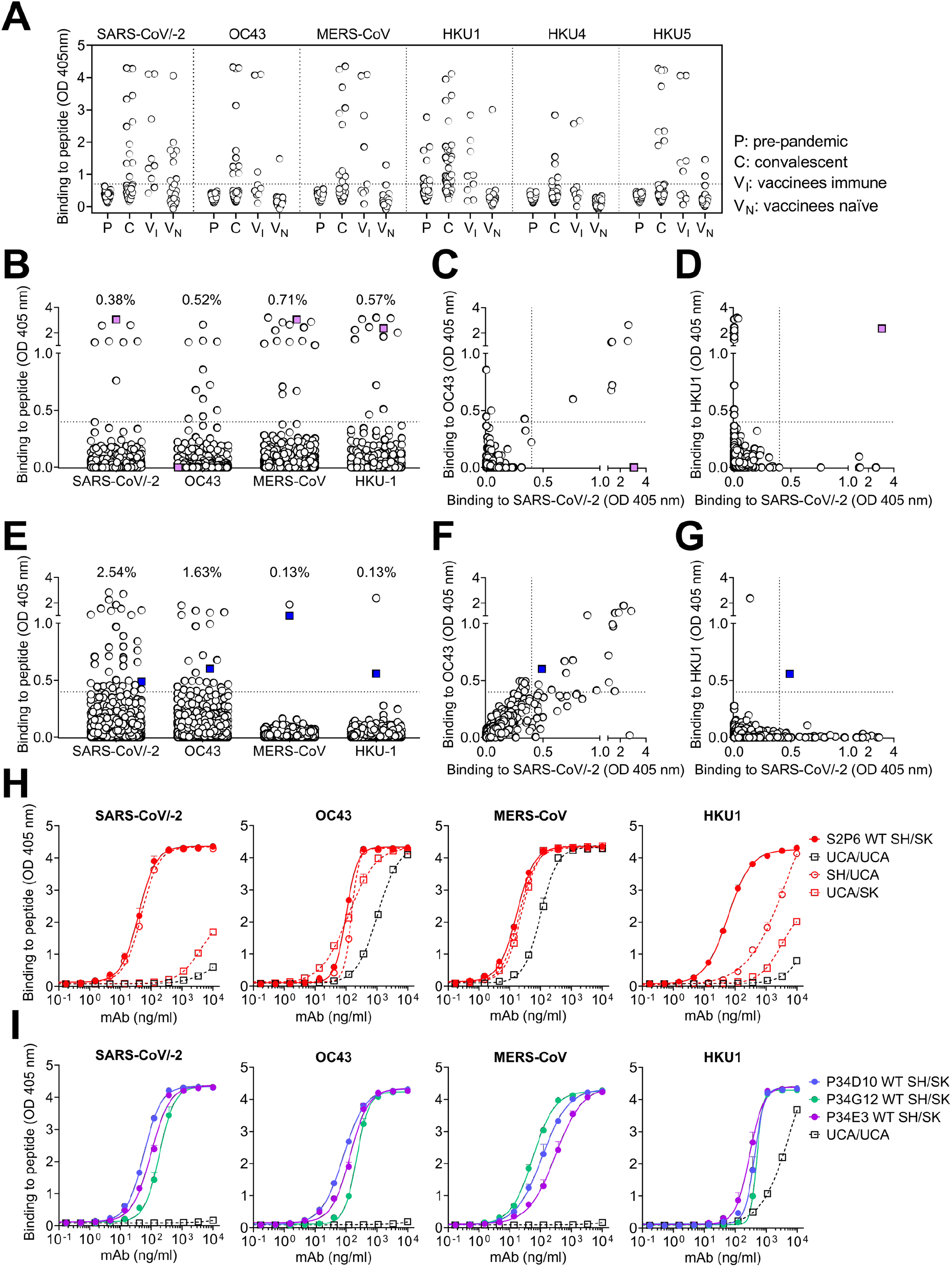
Stem helix-directed Abs are of narrow specificities and acquire affinity and breadth through somatic mutations. (**A**) Binding of pre-pandemic (P, n=88), COVID-19 convalescent (C, n=72), vaccinees immune (VI, n=9) and vaccinees naïve (VN, n=37) plasma Abs diluted 1:10 to β-coronavirus stem helix peptides analyzed by ELISA. A cut-off of 0.7 was determined based on signal of pre-pandemic samples and binding to uncoated ELISA plates. (**B-G**) Analysis of memory B-cell binding to β-coronavirus stem helix peptides from 21 COVID-19 convalescent individuals (B-D) and 16 vaccinees (E-G). Each dot representing individual culture containing oligo-clonal B cells screened against stem helix peptides in ELISA. Pairwise reactivity comparison is shown for SARS-CoV/-2 and OC43 (C and F) and SARS-CoV/-2 and HKU1 (D and G). Highlighted in color are cultures cross-reactive with at least three peptides. (**H-I**) Binding of S2P6 (H) (mutated SH/SK), fully germline reverted (UCA/UCA), germline reverted heavy chain paired with mature light chain (UCA/SK), mature heavy chain paired with germline reverted light chain (SH/UCA) and of P34D10, P34G12 and P34E3 (I) (WT SH/SK), and germline reverted (UCA/UCA) to stem helix peptides.

Next, we investigated the frequency of stem helix specific Abs in the memory B cell repertoire of 21 convalescent and 17 vaccinated individuals using a clonal analysis based on in vitro polyclonal stimulation (*38*), here referred as antigen-specific B-cell memory repertoire analysis (AMBRA) (**Fig. 5B-G** and **fig. S6–S8**). In both cohorts, we observed frequencies of stem helix specific IgGs ranging from 0.1-2.5%, except for one individual (infected and vaccinated with a single dose of mRNA vaccine) for whom we measured an exceptionally high frequency of SARS-CoV/-2 stem helix-specific Abs (**fig. S8**). Most SARS-CoV-2 stem helix specific Abs were found to be cross-reactive with OC43, consistent with the high sequence identity of the stem helices of these two viruses (**Fig. 1H**). Abs specific for the HKU1 S stem helix were found in some individuals but they were not cross-reactive with other β-coronaviruses (**Fig. 5D** and **G**). This analysis revealed a single example of cross-reactivity to all stem helix β-coronavirus peptides tested (**Fig. 5E**), whereas most others Abs show a more limited cross-reactivity among β-coronaviruses.

### Broadly reactive β-coronavirus Abs acquire affinity and breadth through somatic mutations

To define the ontogeny of the broadly reactive β-coronavirus mAbs described here, we generated a panel of germline variants of S2P6, P34D10, P34E3 and P34G12 mAbs. Two out of seven S2P6 heavy chain residues that are mutated relative to germline contribute to epitope recognition (Q32 and H57) whereas none of the 5 light chain mutated residues participate in S binding (**Fig. 2C-D**). To address the role of VH and VK somatic mutations we therefore generated a panel of S2P6 germline variants for the heavy or the light chain, or both variable regions (VH and VK). The fully germline S2P6 (UCA) bound to OC43 and MERS-CoV stem helix peptides (with approximately 1 order of magnitude higher EC50 compared to the mutated mAb) but not to SARS-CoV/-2 or HKU1 peptides (**Fig. 5H**). Somatic mutations in VH were sufficient for high avidity binding to SARS-CoV/-2, whereas both VH and VK mutations were required for optimal binding to HKU1. The presence of residue G103 in CDRH3 was found to be essential for binding to all β-coronaviruses (**fig. S9A**). Collectively, these findings indicate that the S2P6 mAb likely arose in response to OC43 infection, and its specificity was broadened towards SARS-CoV-2 and HKU1 through somatic mutations selected upon natural infection with one or both of these β-coronaviruses. In contrast, analysis of UCA binding of the clonally related P34D10, P34G12 and P34E3 mAbs suggest they were likely primed by HKU1 infection, rather than OC43, and acquired breadth towards the other human β-coronaviruses primarily through somatic mutations in VH (**Fig. 5I** and **fig. S9B**). Together with the serological and B cell repertoire analysis, these findings demonstrate that broadly reactive β-coronavirus Abs may result from priming of virus-specific B cells gaining affinity and breadth through somatic mutations in response to heterotypic coronavirus exposures.

## Discussion

The coronavirus S_2_ subunit (fusion machinery) contains several important antigenic sites, including the fusion peptide and the heptad-repeat 2 regions, and is more conserved than the S_1_ subunit (*39–43*). As a result, it is an attractive target for broad coronavirus detection and neutralization (*44*). The recent identification of 4 cross-reactive mAbs targeting the stem helix unveiled this previously unknown S_2_ subunit epitope, which is conserved among β-coronavirus S glycoproteins (*12–14, 45*), although none of them inhibit members of all three β-coronavirus subgenera (lineages). Here, we identified five mAbs targeting overlapping epitopes in the S stem helix and cross-reacting with human and animal β-coronaviruses. We showed that S2P6 broadly neutralizes all sarbecoviruses, merbecoviruses and embecoviruses evaluated through inhibition of membrane fusion. The exceptionally broad cross-reactivity and neutralization breadth of S2P6 is explained by the conserved nature of the stem helix among β-coronaviruses. Fewer than ~0.06% SARS-CoV-2 sequences have been reported to be mutated between S residues 1146 and 1159 out of more than 1.3 million genomes deposited in GISAID as of April 2021 and no SARS-CoV-2 VOC harbor residue substitutions within this region.

We also provide evidence that a S_2_ subunit-directed mAb protects hamsters from SARS-CoV-2 challenge, including with the SARS-CoV-2 B.1.351 VOC, with a beneficial effect of Fc-mediated effector functions. These data extend previous studies describing the participation of effector functions to in vivo efficacy of SARS-CoV-2 RBD-specific mAbs (*32, 35*) as well as influenza A hemagglutinin stem-specific broadly neutralizing mAbs (*46, 47*). These observations is reminiscent of similar findings for mAbs targeting a highly conserved epitope on the influenza virus hemagglutinin stem, and indicate that the combination of mAb cross reactivity with effector functions may provide particular potency against these viruses (*15*).

The stem helix is presumed to form a 3-helix bundle in prefusion SARS-CoV-2 S and dynamic conformational changes are likely required to expose the otherwise buried hydrophobic epitope, which is surrounded by conserved glycans potentially further shielding this conserved site. We provide evidence that stem helix-targeted Abs are elicited upon natural infection by endemic (OC43 or HKU1) or pandemic (SARS-CoV-2) coronaviruses as well as by COVID-19 mRNA vaccines. However, stem helix-specific Abs are present at low titers in plasma samples of convalescent or vaccinated individuals and at low frequency in their memory B cell repertoire possibly as a result of limited epitope exposure, similarly to the sub-dominance of Abs to the conserved hemagglutinin stem region of influenza A viruses (*46, 48*).

Stem helix-targeted Abs are predominantly of narrow specificities and only few of them mediate broad β-coronavirus neutralization and protection through accumulation of somatic mutations. These findings along with the moderate neutralization potency and low frequency of such mAbs in COVID-19 convalescent or vaccinated subjects indicate that eliciting high enough titers of stem helix-targeted mAbs through vaccination will be a key challenge to overcome to develop pan-β-coronavirus vaccines. We propose that harnessing recent advances in computational protein design, such as epitope-focused vaccine design approaches (*49–51*) and multivalent display (*52–57*), to target the stem helix or the fusion peptide regions might prove necessary to elicit broad β-coronavirus immunity.

## Acknowledgements

We thank Jay C. Nix for x-ray data collection, and Tristan I. Croll for assistance in refinement of the crystal structure. Thanks to Ann Arvin for her insightful comments. We thank Marcel Meury for help with protein production. We thank Hideki Tani (University of Toyama) for providing the reagents necessary for preparing VSV pseudotyped viruses. The authors would also like to thank Cindy Castado and Normand Blais (GSK Vaccines) for their help in the selection of the genetically divergent sarbecoviruses used in this study. We thank Promega Corporation for kindly providing SARS-CoV2 CHO-K1 cells (genetically engineered to stably express a HaloTag-HiBit-tagged). This study was supported by the National Institute of Allergy and Infectious Diseases (DP1AI158186 and HHSN272201700059C to D.V., and U01 AI151698-01 to WCVV), a Pew Biomedical Scholars Award (D.V.), Investigators in the Pathogenesis of Infectious Disease Awards from the Burroughs Wellcome Fund (D.V.), Fast Grants (D.V.), the Swiss National Science Foundation (P400PB_183942 to M.M.S.), the University of Washington Arnold and Mabel Beckman cryoEM center, the Helmut Horten Foundation (F.S. and the Institute for Research in Biomedicine). The project was also partially funded by the Swiss Kidney Foundation. Use of the Stanford Synchrotron Radiation Lightsource, SLAC National Accelerator Laboratory, is supported by the U.S. Department of Energy, Office of Science, Office of Basic Energy Sciences under Contract No. DE-AC02-76SF00515. The SSRL Structural Molecular Biology Program is supported by the DOE Office of Biological and Environmental Research, and by the National Institutes of Health, National Institute of General Medical Sciences (P30GM133894). The contents of this publication are solely the responsibility of the authors and do not necessarily represent the official views of NIGMS or NIH.

## Author contributions

Experiment design: D.P., M.M.S., J.S.L., M.A.T., M.S.P., H.W.V, MB., D.C., D.V. Donors’ Recruitment and Sample Collection: E.C., F.B., B.G., A.C., P.F., P.E.C., O.G.,_S.C., C.G., A.R., L.P., M.S., D.V., C.H.D. Antibody discovery: D.P., MB., J.S.L., J.J., S.J., N.S., K.C., E.C., D.C. AMBRA preparation: C.S.F., J.B., A.C. Expression and purification of proteins: A.C.W., J.E.B., J.J. Antibody functional experiments: D.P., M.M.S., N.C., J.S.L., M.A.T., A.C.W., F.L., J.N., S.B., R.M. Bioinformatic analysis of virus diversity and variants: J.d.I., I.B., A.T. Evaluation of effector functions: B.G. SPR Binding Assays: L.E.R. Neutralization assay: D.P., M.A.T., F.A.L., J.N., M. P. H. Escape mutants selection and sequencing: H.K., S.I., F.A.L., J.d.I. Effects in the hamster model and data analysis: R.A., C.F., C.D.K., L.B, L..C, J.N., E.V., F.B. Cryo-EM Data Collection, Processing, and Model Building, M.M.S., D.V. Crystallization, X-Ray Crystallography Data Collection, Processing, and Model Building: M.M.S., N.C., G.S., D.V. Serological Assays: D.P., R.M. Data Analysis: D.P., M.M.S., M.B., J.S.L., J.J., F.A.L., J.N., N.C., A.M.T., A.C.W., G.S., D.C., D.V. Manuscript Writing: D.O.P., M.M.S., J.S.L., M.B., G.S., F.S., A.L., H.W.V, D.C., D.V.

## Competing interests

D.P., N.C., M.P.H., J.N., B.G., L.E.R., J.d.I., H.K., S.I., S.J., N.S., K.C., I.B., S.B., C.S.F, J.B., R.M., E.V., F.B., E.C., L.P., M.S.P., M.S., D.H., A.T., F.A.L., C.H.D., A.L., G.S., H.W.V., M.B. and D.C. are employees of Vir Biotechnology and may hold shares in Vir Biotechnology. D.C., J.S.L, F.S., A.C. and A.L. are currently listed as an inventor on multiple patent applications, which disclose the subject matter described in this manuscript. D.V. is a consultant for Vir Biotechnology Inc. The Veesler laboratory and the Sallusto laboratory have received sponsored research agreements from Vir Biotechnology Inc. The other authors declare no competing interests.

## Materials and Methods

### Cell lines

Cell lines used in this study were obtained from ATCC (HEK293T and Vero-E6) or ThermoFisher Scientific (ExpiCHO cells, FreeStyle™ 293-F cells and Expi293F™ cells).

### Sample donors

Samples were obtained from cohorts of individuals enrolled before June 2019 (pre-pandemic), of SARS-CoV-2 infected individuals or of vaccinated individuals immunized with Moderna or Pfizer/BioNTech BNT162b2 vaccines under study protocols approved by the local Institutional Review Boards (Canton Ticino Ethics Committee, Switzerland, the Ethical committee of Luigi Sacco Hospital, Milan, Italy and WCG North America, Princeton, NJ, US). All donors provided written informed consent for the use of blood and blood components (such as human peripheral blood mononuclear cells (PBMCs), sera or plasma) and were recruited at hospitals or as outpatients. PBMCs were isolated from blood by Ficoll density gradient centrifugation and either used freshly or stored in liquid nitrogen for later use. Sera were obtained from blood collected using tubes containing clot activator, followed by centrifugation and stored at −80°C.

### AMBRA (antigen-specific memory B cell repertoire analysis) of IgG antibodies

Replicate cultures of total unfractionated PBMC from SARS-CoV-2 infected or vaccinated individuals were seeded in 96 U-bottom plates (Corning) in RPMI1640 supplemented with 10% Hyclone, sodium pyruvate, MEM non-essential amino acid, stable glutamine and PenicillinStreptomycin. Memory B cell stimulation and differentiation was induced by adding 2.5 μg/ml R848 (3 M) and 1000 U/ ml human recombinant IL-2 for 10 days at 37 °C 5% CO_2_. The cell culture supernatants were collected for further analysis.

### Antibody discovery and expression

Antigen specific IgG^+^ memory B cells were isolated and cloned from total PBMCs of convalescent individuals. Abs VH and VL sequences were obtained by reverse transcription PCR (RT-PCR) and mAbs were expressed as recombinant human IgG1, carrying the half-life extending M428L/N434S (LS) mutation in the Fc region or Fab fragment. ExpiCHO cells were transiently transfected with heavy and light chain expression vectors as previously described (*21*). For in vivo experiments in Syrian hamsters, S2P6 was produced with a Syrian hamster IgG2 constant region. Using the Database IMGT (http://www.imgt.org), the VH and VL gene family and the number of somatic mutations were determined by analyzing the homology of the VH and VL sequences to known human V, D and J genes. UCA sequences of the VH and VL were constructed using IMGT/V-QUEST.

MAbs affinity purification was performed on ÄKTA Xpress FPLC (Cytiva) operated by UNICORN software version 5.11 (Build 407) using HiTrap Protein A columns (Cytiva) for full length human and hamster mAbs and CaptureSelect CH1-XL MiniChrom columns (ThermoFisher Scientific) for Fab fragments, using PBS as mobile phase. Buffer exchange to the appropriate formulation buffer was performed with a HiTrap Fast desalting column (Cytiva). The final products were sterilized by filtration through 0.22 μm filters and stored at 4°C.

### Flow cytometry of antibody on S Protein expressing ExpiCHO-S cells

For Expi-CHO cell transient transfection, S plasmids (*21, 58*) were diluted in cold OptiPRO SFM, mixed with ExpiFectamine CHO Reagent (Life Technologies, A29130) and added to the cells seeded at 6×10^6^ cells/ml in a volume of 5 ml in a 50 ml bioreactor. Transfected cells were incubated at 37°C, 8% CO_2_ with an orbital shaking speed of 209 rpm (orbital diameter of 25 mm) for 42 hours. To test mAb binding, transfected ExpiCHO cells were collected, washed twice in wash buffer (1% w/v solution of Bovine Serum Albumin (BSA; Sigma) in PBS, 2 mM EDTA) and distributed at 60,000 cells/well into 96 U-bottom plates (Corning). mAb serial dilutions from 10 μg/ml were added onto cells for 30 minutes on ice and, after two washes, Alexa Fluor647-labelled Goat Anti-Human IgG (Jackson Immunoresearch, 109-606-098) was used for detection. After 15 minutes of incubation on ice, cells were washed twice and mAb binding analyzed by flow cytometry using a ZE5 Cell Analyzer (Biorard).

### Protein expression and purification

SARS-CoV-2 S 2P, SARS-CoV S 2P, MERS-CoV S 2P, OC43 S, HKU1 S2P, and HKU4 S2P ectodomains were produced as previously described (*3, 11, 12, 59, 60*). SARS-CoV-2 S D614G, used for production of SARS-CoV-2 postfusion, contains a mu-phosphatase signal peptide beginning at 14Q, a mutated S1/S2 cleavage site (SGAR), and ends at residue K1211 followed by a TEV cleavage, foldon trimerization motif, and an 8X his tag in a pCMV vector. Briefly, spike glycoproteins were produced in Expi293F cells grown in suspension using Expi293 expression medium (Life Technologies) at 37°C in a humidified 8% CO_2_ incubator rotating at 130 rpm. The cultures were transiently transfected using PEI with cells grown to a density of 3 million cells per mL and cultivated for 3 days. The supernatant was clarified and affinity purified using a 1 mL HisTrapFF column (Cytiva). To isolate post-fusion SARS-CoV-2 S, SARS-CoV-2 S D614G ectodomain was incubated for one hour with the S2X58 triggering Fab (*61*) and 1 μg/ml trypsin before size-exclusion chromatography purification using a Superose 6 Increase 10/24 column (Cytiva). Purified protein was concentrated, quantified using absorption at 280 nm, and flash frozen in Tris-saline (20 mM Tris pH 8.0, 100 mM NaCl).

### Enzyme-linked immunosorbent assay (ELISA)

96-well plates (Corning) were coated overnight at 4°C with recombinant proteins at 1 μg/ml or peptides at 8 μg/ml diluted in phosphate-buffered saline (PBS). Plates were blocked with a 1% w/v solution of Bovine Serum Albumin (BSA; Sigma) in PBS and serial dilutions of mAbs were added for 1 hour at room temperature. When testing human plasma or memory B-cell supernatants, plates were blocked with Blocker Casein (1% w/v) in PBS (Thermo Fisher Scientific) supplemented with 0.05% Tween 20. Plasma and memory B-cell supernatants (AMBRA testing) were then incubated for 1 hour at room temperature at a 1:10 and 1:2 dilution, respectively. After further wash, mAbs bound were revealed using an anti-human IgG coupled to alkaline phosphatase (Jackson Immunoresearch) incubated for 1 hour. Substrate (p-NPP, Sigma) was used for color development and plates read at 405 nm by a microplate reader (Biotek). The data were plotted with GraphPad Prism software.

For ELISA with plasma, cut-off value (OD=0.7) was determined based on signal of pre-pandemic samples and binding to uncoated ELISA plates. For AMBRA, cut off value (OD= 0.4) was determined as three times the mean OD values of negative wells.

### Blockade of SARS-CoV-2 S binding to ACE2

SARS-CoV-2 S prefusion (final concentration 300 ng/ml) was incubated with 1 μg/ml of S309 mouse Fc-tagged mAb (S309-mFc) 30 minutes at 37°C before the addition of serially diluted S2P6 (from 20 μg/ml) and incubated for additional 30 minutes at 37°C. The complex S:S309:S2P6 was then added to a pre-coated hACE2 (2 μg/ml in PBS) 96-well plate MaxiSorp (Nunc) and incubated 1 hour at room temperature. Subsequently, the plates were washed and a goat anti-mouse IgG (Southern Biotech) coupled to alkaline phosphatase (Jackson Immunoresearch) added to detect SARS-CoV-2 S:S309-mFc binding. After further washing, the substrate (p-NPP, Sigma) was added, and plates read at 405 nm using a microplate reader (Biotek). The percentage of inhibition was calculated as follow: (1-((OD sample-OD neg ctr)/(OD pos. ctr-OD neg. ctr))*100.

### Epitope identification and substitution scan

PEPperMAP Epitope Mapping (PEPperPRINT GmbH, Heidelberg, Germany) was performed to determine mAbs epitope through a pan-corona Spike protein Microarray covering the S proteins of all β-coronaviruses. Briefly, microarray containing 15-mer peptides (overlapping of 13-mer) was incubated with 10 μg/ml mAb for 16 hours at 4°C shaking at 140 rpm followed by staining with Goat anti-human IgG (H+L) DyLight680 for 45 minutes at room temperature. Microarray read-out was performed with a LI-COR Odyssey Imaging System at scanning intensities of 7/7 (red/green). Epitope substitution scan was performed on the identified epitope based on a stepwise single amino acid exchange on all amino acid positions. The mAbs binding to the generated microarray was performed as above.

### Conservation analysis

Conservation analysis was performed as described previously (Pinto et al 2020). SARS-CoV-2 S sequences were obtained from GISAID (https://www.gisaid.org/) on Apr 22^nd^ 2021, the other viruses sequences were obtained from NCBI Virus (https://www.ncbi.nlm.nih.gov/labs/virus/vssi/#/) in December 2020. The multiple sequences alignment was performed using MAFFT (https://mafft.cbrc.jp/alignment/software/) with the spike amino acid sequences as input.

### SPR binding measurements

SPR binding measurements were performed using a Biacore T200 instrument using anti-AviTag pAb covalently immobilized on CM5 chips to capture S ECDs except the Cytiva Biotin CAPture kit was used to capture biotinylated OC43 S ECD. Running buffer was Cytiva HBS-EP+ (pH 7.4) or 20 mM phosphate pH 5.4, 150 mM NaCl, 0.05% P-20, for neutral or acidic pH experiments, respectively. All measurements were performed at 25 °C. S2P6 Fab or IgG concentrations were 11, 33, 100, and 300 nM run as single-cycle kinetics. Double reference-subtracted data were fit to a binding model using Biacore Evaluation software. All data for SARS-CoV-2 S, SARS-CoV S, and OC43 S were fit to a 1:1 binding model. Data for MERS S were fit to a Heterogeneous Ligand binding model, due to a kinetic phase with very slow dissociation presumed to be an artifact; the lower affinity of the two KDs returned by the fit is reported as the KD of the S2P6:MERS S interaction and is indicated to be approximate (the Rmax associated with the higher affinity kinetic phase is proportional to the magnitude of the final signal above baseline). Data for HKU1 S were fit to a steady-state binding model, because of the low signal and fast approach to equilibrium within each association phase; the reported KD is indicated to be approximate. IgG binding data yield an “apparent KD” due to avidity.

### Neutralization of authentic SARS-CoV-2 virus

For SARS-CoV-2 neutralization experiments, cells were cultured in DMEM (Gibco 11995-040) supplemented with 10% FBS (VWR 97068-085 lot#345K19) and 100 U/ml Penicillin-Streptomycin (Gibco 15140-122). Cells were seeded in black, 96-well glass bottom plates (Cellvis P96-1.5H-N) at a density of 20,000 cells/well. In a BSL3 facility, serial dilutions of mAbs (1:4) were incubated with 200 PFU (plaque forming units, corresponding to a multiplicity of infection of 0.01) of authentic SARS-CoV-2 (isolate USA-WA1/2020, passage 3, passaged in Vero-E6 cells) for 30 minutes at 37°C. After removal of cell culture supernatants, cells were infected with the virus:mAb mixtures and incubated for 20 hours at 37°C. Cells were then fixed with 4% paraformaldehyde (Electron Microscopy Sciences, 15714-S) in PBS (Gibco 10010-031) for 30 minutes, permeabilized with 0.1% Triton X-100 (Sigma, X100-500ML) for 30 minutes, and stained with Human SARS Coronavirus Nucleoprotein/NP Ab, Rabbit Mab (Sino Biological, 40143-R001) at a dilution of 1:2000 dilution in 2% milk (RPI, M17200-500.0) for 1 hour. Subsequently, cells were stained with Goat anti-Rabbit IgG (H+L) AF647 (Invitrogen, Cat. A21245 Lot. 223 2862) at a dilution of 1:1000 and 2 ug/ml Hoechst 33342 in 2% milk for 1 hour. Plates were imaged with an automated microscope (Cytation5, Biotek), and nuclei and cells positive for the SARS-CoV-2 Nucleoprotein were quantified using the supplied Gen5 software.

### VSV pseudotype virus production and neutralization

Sarbecovirus spike cassettes with a C-terminal deletion of 19 amino acids (D19) were synthesized and cloned into mammalian expression constructs (pcDNA3.1(+) or pTwist-CMV) for the following Sarbecoviruses: SARS-CoV-2 (Accession QOU99296.1), SARS-CoV-1 (Accession AAP13441.1), hCoV-19/pangolin/Guangdong/1/2019 (GD19, Accession QLR06867.1), and Middle East respiratory syndrome-related coronavirus (MERS, Accession YP_009047204). To generate pseudotyped VSV, 293T Lenti-X packaging cells (Takara, 632180) were seeded in 15 cm dishes such that the cells would be 80% confluent the following day. Cultures were then transfected with various S expression plasmids using TransIT-Lenti transfection reagent (Mirus, 6600) according to the manufacturer’s instructions. 24 hours after transfection, the packaging cells were infected with VSV-G*ΔG-luciferase (Kerafast, EH1020-PM). 48 hours after infection the supernatant containing Sarbecovirus pseudotyped VSV-luc was collected, centrifuged at 1000 × g for 5 minutes, aliquoted and frozen at −80°C.

To perform pseudotype neutralization assays, VeroE6-TMPRSS2 cells were used for VSV-SARS-CoV-2, VSV-SARS-CoV-1, and VSV-GD19 and Huh7 cells were used for VSV-MERS. Cells were seeded into clear bottom white-walled 96-well plates at 20,000 cells/well. The following day, 1:3 serial dilutions of Ab were prepared in DMEM and pseudotyped VSVs (final dilution 1:20) were added to each mAb dilution and incubated for 1 hour at 37°C. Media was removed from the cells and replaced with 50 μl of pseudotype:mAb complex and one hour post-infection, 50 μl of complete culture media was added to the cells and incubated overnight at 37°C. The media from infected cells was then removed and 100 μl of 1:1 diluted PBS:Bio-Glo (Promega, G7940) luciferase substrate was added to each well. The plates were shaken at 300 rpm at room temperature for 10 minutes and relative light units (RLUs) were then read on an EnSight microplate reader (Perkin Elmer). Percent neutralization was determined by subtracting the mean background (uninfected cells with luciferase substrate alone) values of 6 wells per plate from all data points. Percent neutralization for each mAb concentration was calculated relative to control wells receiving no mAb for each plate. Percent neutralization data were analyzed using GraphPad Prism. Absolute IC50 values were calculated by fitting a curve using a variable slope 4 parameter non-linear regression model and values were interpolated from the curve at y=50.

Production of OC43 S (AAT84354.1) pseudotyped VSV virus and neutralization assays was performed similarly to previously described (*12*). Briefly, HEK-293T cells at 70~80% confluency were transfected with the pCDNA3.1 expression vectors encoding full-length OC43 S harboring a truncation of the 17 C-terminal residues along with a fusion to Ha-tag and the bovine coronavirus hemagglutinin esterase protein Fc-tagged at molar ratios of 7:1. The day after, cells were transduced with VSVΔG/Fluc (*20*). After 2 h, infected cells were washed four times with DMEM before adding medium supplemented with anti-VSV-G antibody (I1-mouse hybridoma supernatant diluted 1 to 25, from CRL-2700, ATCC). Supernatant was harvested 18-24 h post-inoculation, clarified from cellular debris by centrifugation at 2,000 x g for 5 min and concentrated 10 times using a 30 kDa cut off membrane and aliquoted and frozen at −80°C until use in neutralization experiments. For viral neutralization, stable HRT-18G cells (ATCC) in DMEM supplemented with 10% FBS, 1% PenStrep were seeded at 40,000 cells/well into clear bottom white walled 96-well plates and cultured overnight at 37°C. Twelve-point 3-fold serial dilutions of S2P6 were prepared in DMEM and OC43 S VSV pseudoviruses were added 1:1 (v/v) to each dilution in the presence of anti-VSV-G antibody from I1-mouse hybridoma supernatant diluted 50 times (final volume: 50 μl). After 45 min incubation at 37 °C, 40 μl of the mixture was added to the cells and 2 h post-infection, another 40 μL DMEM were added to avoid evaporation. After 17-20 h, 50 μL/well of One-Glo-EX substrate (Promega) were added to the cells and incubated in the dark for 5-10 min prior reading on a Varioskan LUX plate reader (ThermoFisher). Data was processed using GraphPad Prism v9.0.

### Selection of VSV-SARS-CoV-2 mAb escape mutants

#### Resistant virus selection

Cells were cultured in DMEM (Gibco 11995-040) supplemented with 10% FBS (VWR 97068-085 lot#345K19) and 100 U/ml Penicillin-Streptomycin (Gibco 15140-122). The day before infection, 250,000 VeroE6-TMPRSS2 cells were seeded in 12-well plates in 2 ml of DMEM (Gibco 11995-040) supplemented with 10% FBS (VWR 97068-085 lot#345K19) and 100 U/ml Penicillin-Streptomycin (Gibco 15140-122) and incubated overnight at 37C. The next day, S2P6 was serially diluted 1:4 starting at 80 μg/ml in infection media (DMEM supplemented with 2% FBS and 20mM HEPES (Gibco, 15630-080)) and incubated with replication-competent VSV-SARS-CoV-2 (*23*) at MOI 2 for 1 hour at 37°C. A no Ab control was included to account for any tissue culture adaptations and quasispecies variability that may occur during virus replication. The mAb-virus complexes were adsorbed on the cells for 1 hour at 37°C, with manual rocking every 15 minutes. After adsorption, cells were washed with PBS and overlaid with infection media containing an equivalent amount of S2P6 as was used for the initial infection. Infection was monitored visually by microscopy for GFP expression and cytopathic effect (CPE) of the cells at day 1 and day 3 post-infection. At day 3 post-infection, when the no mAb control well reached >50% CPE, the well with the highest Ab concentration showing >20% CPE (in this case the 80 ug/ml well) was selected for passaging. The cell supernatant was centrifuged to remove cell debris, diluted 1:10 in infection medium and added to fresh VeroE6-TMPRRS2 cells with the same S2P6 concentration range and treatment as for the initial passage. Selection was stopped after two passages, after no virus neutralization was observed at the highest concentration tested.

#### Sequencing of S gene

Viral RNA was extracted from the supernatant of viral passages using the QIAamp Viral RNA Mini Kit (Qiagen, 52904) according to the manufacturer’s instructions, without the addition of carrier RNA. Reverse transcription reactions were performed with 6 μl of purified RNA and random primers using the NEB ProtoScript II First Strand cDNA Synthesis Kit (NEB, E6560S), according to manufacturer’s instructions. The resulting cDNA was used as a template for PCR amplification of the spike gene using the KapaBiosystems polymerase (KAPA HiFi HotStart Ready PCR Kit KK2601) with primers 5’-CGAGAAAAAGGCATCTGGAG-3’ and 5’-CATTGAACTCGTCGGTCTC-3’. Amplification conditions included an initial 3 minutes at 95°C, followed by 28 cycles with 20 seconds at 98°C, 15 seconds at 59°C and 72°C for 2 minutes, with a final 4 minutes at 72°C. PCR products were purified using AMPure XP beads (Beckman Coulter, A63881) following manufacturer’s instructions. The size of the amplicon was confirmed by analyzing 2 μl of PCR products using the Agilent D5000 ScreenTape System (Agilent D5000 ScreenTape, 5067-5588, Agilent D5000, Reagents 5067-5589). Products were quantified by analyzing 2 μl with the Quant-iT dsDNA High-Sensitivity Assay Kit (Thermo Fisher, Q331120). Twenty ng of purified PCR product was used as input for library construction using the NEBNext Ultra II FS DNA Library Prep Kit (NEB, E6177S) following manufacturer’s instructions. DNA fragmentation was performed for 13 minutes. NEBNext Multiplex Oligos for Illumina Dual Index Primer Set 1 (NEB, E7600S) was used for library construction, with a total of 6 PCR cycles. Libraries size was determined using the Agilent D1000 ScreenTape System (Agilent D1000 ScreenTape, 5067-5582, Agilent D5000 Reagents, 5067-5583) and quantified with the Quant-iT dsDNA High-Sensitivity Assay Kit. Equal amounts of each library were pooled together for multiplexing and ‘Protocol A: Standard Normalization Method’ of the Illumina library preparation guide was used to prepare 8 pM final multiplexed library with 1% PhiX spike-in for sequencing. The Illumina MiSeq Reagent Kit v3 (600-cycle) (Illumina, MS-102-300) was used for sequencing the libraries on the Illumina MiSeq platform, with 300 cycles for Read 1,300 cycles for Read 2, 8 cycles for Index 1, and 8 cycles for Index 2.

#### Bioinformatic analysis

The average read length after running Illumina’s Bcl2fastq command was ranging from 149 to 188bp on average per sample. For consistency across samples, paired-end reads were initially trimmed to 2X150bp and further cleaned to remove Illumina’s adapter and low quality bases using Trimmomatic (*62*). Read alignment was performed with Burrows-Wheeler Aligner (BWA (*63*)) using a custom reference sequence. Variants were called with LoFreq upon indel realignment and base quality recalibration (*64*), using a frequency threshold of 1%. Two consecutive rounds of alignments and variant calling were performed, where the variants called during the first round at allelic frequency >50% were integrated in the reference for the second round in order to adjust alignment rate and variant calling accuracy. Variants were annotated with SnpEff (*65*). The reference sequence coordinates were mapped back to the SARS-CoV-2 Wuhan-Hu-1 sequence (NCBI: NC_045512.2) in order to match the reference sequence nomenclature. Extensive QCs were performed at read, alignment and variant level using FastQC, samtools, picard, mosdepth (*66*), bcftools_ (*67*), MultiQC (*68*) and in-house scripts, notably to remove variants that were consistently called at a static position in reads (such as the beginning or end of reads that were carrying it, rather than being randomly distributed throughout those reads.). An end-to-end workflow was automated using NextFlow (*69*). All programs are available through the Bioconda Initiative (*70*)(bioconda.github.io).

### Crystallization and structure determination

Crystals of the S2P6 Fab/SARS-CoV peptide complex were obtained using the sitting-drop vapor diffusion method at 20°C with a Fab concentration of 12 mg/ml and a 1.5x molar excess of peptide. A total of 150 nl S2P6 Fab/peptide solution in 20 mM Tris-HCl pH 7.5, 50 mM NaCl were mixed with 150 nl mother liquor containing 0.2 M ammonium sulfate, 0.1 M sodium acetate pH 4.6 and 25% (v/v) PEG Smear Broad (Molecular Dimensions). Crystals were flash frozen in liquid nitrogen. Data were collected at beamline 12-2 at the Stanford Synchrotron Radiation Lightsource facility in Stanford, CA. Data were processed with the XDS software package (Kabsch, 2010) for a final dataset of 2.67 Å in space group P6_5_22. The S2P6 Fab/peptide complex structure was solved by molecular replacement using a homology model of the S2P6 Fab built using the Molecular Operating Environment (MOE) software package from the Chemical Computing Group (https://www.chemcomp.com). Several subsequent rounds of model building and refinement were performed using Coot (*71*), ISOLDE (*72*), Refmac5 (*73*), and MOE, to arrive at a final model for the complex.

### Measurement of Fc-effector functions

#### MAb-dependent activation of human FcγRIIIa and FcγRIIa

Determination of mAb-dependent activation of human FcγRIIIa and FcγRIIa was performed using ExpiCHO cells stably expressing full-length wild-type SARS-CoV-2 spike (S) (target cells). Cells were incubated with different amounts of mAbs for 10 minutes before incubation with Jurkat cells stably expressing FcγRIIIa receptor (V158 variant) or FcγRIIa receptor (H131 variant) and NFAT-driven luciferase gene (effector cells) at an effector to target ratio of 6:1 for FcγRIIIa and 5:1 for FcγRIIa. Activation of human FcγRs was quantified by the luciferase signal produced as a result of NFAT pathway activation. Luminescence was measured after 21 hours of incubation at 37°C with 5% CO_2_ with a luminometer using the Bio-Glo-TM Luciferase Assay Reagent according to the manufacturer’s instructions (Promega, Cat. Nr.: G7018 and G9995).

#### Antibody-dependent cell cytotoxicity (ADCC)

ADCC assays were performed using SARS-CoV2 CHO-K1 cells (genetically engineered to stably express a HaloTag-HiBit-tagged) as target cells and PBMC as effector cells at a E:T ratio of 33:1. HiBit-cells were seeded at 3,000 cells/well and incubated for 16 hours at 37°C, while PBMCs isolated from fresh blood (VV donor) were cultivated overnight at 37°C 5% CO_2_ in the presence of 5 ng/ml of IL-2. The day after, media was removed and titrated concentrations of mAbs were added before the addition of PBMCs at 100,000 cells/well. As 100% specific lysis, Digitonin at 100 ug/ml was used. After 4 hours of incubation at 37°C, ADCC was measured with Nano-Glo HiBiT Extracellular Detection System (Promega; Cat. Nr.: N2421) using a luminometer (Integration Time 00:30).

#### Antibody-dependent cellular phagocytosis (ADCP)

ADCP was performed using CHO cells stably expressing full-length wild-type SARS-CoV-2 S glycoprotein (target cells) fluorescently labelled with PKH67 Fluorescent Cell Linker Kits (Sigma Aldrich; Cat. Nr.: MINI67). Target cells were incubated with titrated concentrations of mAbs for 10 minutes, followed by incubation with PBMCs fluorescently labelled with Cell Trace Violet (Invitrogen, cat. no. C34557) after an overnight incubation in 5 ng/ml IL-2 (Recombinant Human Interleukin-2; ImmunoTools GmbH; Cat. Nr.: 11340027). An effector:target ratio of 20:1 was used. After an overnight incubation at 37°C, cells were stained with anti-human CD14-APC Ab (BD Pharmingen, cat. no. 561708, Clone M5E2) to stain monocytes. ADCP was determined by flow cytometry, gating on CD14+ cells that were double-positive for cell trace violet and PKH67.

#### Complement-dependent cytotoxicity (CDC)

CDC was performed on CHO cells stably expressing SARS-CoV-2 S glycoprotein (target cells) incubated with serial dilutions of mAbs for 10 minutes, followed by incubation with pre-adsorbed Low-Tox M Rabbit Complement (Cederlane Laboratories Limited; Cat. Nr.: CL3051) at a final dilution of 1:12. CDC was measured after incubation for 3 hours at 37 °C 5% CO_2_ with a luminometer using the CytoTox-Glo Cytotoxicity Assay (Promega; Cat. Nr.: G9291) according to the manufacturer’s instructions.

### S2P6 binding and S2P6/B6 competition experiments to different synthetic coronavirus S stem peptides

All biotinylated coronavirus stem helix peptides binding experiments were performed in PBS supplemented with 0.005 % Tween20 (PBST) at 30°C and 1,000 rpm shaking on an Octet Red instrument (Fortebio). For S2P6 binding to different stem helix peptides, 1 μg/ml biotinylated stem peptide (15- or 16-residue long stem peptide-PEG6-Lys-Biotin synthesized from Genscript) was loaded on SA biosensors to a threshold of 0.5 nm. Then, the system was equilibrated in PBST for 300 seconds prior to immersing the sensors in 0.1 μM S2P6 mAb, respectively, for 300 seconds prior to dissociation in buffer for 300 seconds. For S2P6-B6 competition, 1 μg/ml biotinylated to SARS CoV-2 peptide was loaded on SA biosensors to a threshold of 0.5 nm. The system was equilibrated in PBST for 180 seconds and each subsequent step was monitored for 900 seconds. The first sample biosensor was immersed in 0.1 μM mAb S2P6 prior to immersing the sample biosensor in a solution of 0.1 μM mAb S2P6 and B6, respectively. The second sample biosensor was immersed in PBST and subsequently in 0.1 μM mAb B6. To monitor unspecific binding, identical experiments were performed without loading stem peptides to the biosensors.

### CryoEM sample preparation and data collection

1 mg/ml SARS-CoV-2 S 2P was incubated with 1.5-fold molar excess of S2M11 Fab for 30 minutes at 37°C (to promote the closed trimer conformation). Excess S2M11 Fab was removed from the sample using a centrifugal filter (amicon ultra, 100kDa cut off). Then a 2-fold molar excess of S2P6 Fab over SARS-CoV-2 S protomer was added to the solution and incubated for additional 45 minutes at 37°C. 3 μl sample were applied on to a freshly glow discharged UltrAUFoil Au 200 (R2/2) grid. Plunge freezing was performed using a TFS Vitrobot Mark IV (blot force: 0, blot time: 6.5 s, Humidity: 100 %, temperature: 23°C). Data were acquired using a FEI Titan Krios transmission electron microscope operated at 300 kV and equipped with a Gatan K3 Summit direct detector and Gatan Quantum GIF energy filter, operated in zero-loss mode with a slit width of 20 eV. Automated data collection was carried out using Leginon (*74*) at a nominal magnification of 105,000x with a pixel size of 0.4215Å. The dose rate was adjusted to 15 counts/pixel/s, and each movie was acquired in super-resolution mode fractionated in 75 frames of 40 ms. Tilted data collection (45° tilt) was performed to compensate for preferential specimen orientation and 6,015 micrographs were collected in a single session with a defocus range comprised between 0.5 and 5.0 μm.

### CryoEM data processing

Movie frame alignment, estimation of the microscope contrast-transfer function parameters, particle picking and extraction were carried out using Warp (*75*). Particle images were extracted with a box size of 1024 pixels^2^ binned to 256 pixels^2^ yielding a pixel size of 1.686 Å. Two rounds of reference-free 2D classification were performed using cryoSPARC (*76*) to select well-defined particle images. Subsequently, one round of 3D classification with 25 iterations was carried out using Relion without imposing symmetry and using an initial ab initio model created in cryoSPARC.

The best subclasses were combined and non-uniform refinement (NUR), defocus refinement (DR) and NUR again performed in cryoSPARC. We then performed one round of global CTF refinement of beam-tilt, trefoil and tetrafoil parameters followed by another refinement cycle of NUR-DR-NUR. Selected particle images were then subjected to Bayesian polishing in Relion (*77*). During this step the box and pixel size were changed to 426 pixels and 1.201 Å, respectively, before performing another NUR-DR-NUR refinement cycle. We then performed one additional round of focused classification in Relion with 25 iterations, skipping the oriental assignment and using a mask covering the strongest S2P6 Fab density and a small part of the S stem to further separate distinct S2P6 Fab conformations. The best classes were combined and a final round of NUR performed. Reported resolutions are based on the gold-standard Fourier shell correlation (FSC) of 0.143 criterion and Fourier shell correlation curves were corrected for the effects of soft masking by high-resolution noise substitution (*78*).

### CryoEM model building and analysis

UCSF Chimera (*79*) was used to fit atomic models into the cryoEM maps. The SARS-CoV-2 S EM structure in complex with the variable domain of the S2M11 Fab (PDB 7K43, residue 15-1140), the constant domain of the S2H14 Fab crystal structure and the S2P6-SARS-CoV-2 (residue 1146-1159) crystal structure were fit into the cryoEM map.

### Fusion inhibition assay

For testing inhibition of spike-mediated cell-cell fusion Vero-E6 cells were seeded in 96 well plates at 20,000 cells/ well in 70 μl DMEM with high glucose and 2.4% FBS (Hyclone). After 16 hours, cells were transfected with SARS-CoV-2-S-D19_pcDNA3.1 as follows: for 10 wells, 0.57 μg plasmid SARS-CoV-2-S-D19_pcDNA3.1 were mixed with 1.68 μl X-tremeGENE HP in 30 μl OPTIMEM. After 15 minutes incubation, the mixture was diluted 1:10 in DMEM medium and 30 μl was added per well. A 4-fold serial dilution mAb was prepared and added to the cells, with a starting concentration of 20 μg/ml. The following day, 30 μl 5X concentrated DRAQ5 in DMEM was added per well and incubated for 2 hours at 37°C. Nine images of each well were acquired with a Cytation 5 equipment for analysis.

### In vivo mAb testing using a Syrian hamster model

KU LEUVEN R&D has developed and validated a SARS-CoV-2 Syrian Golden hamster infection model (*36*).

#### SARS-CoV-2 virus production

The wt SARS-CoV-2 strain used in this study, BetaCov/Belgium/GHB-03021/2020 (EPI ISL 109 407976|2020-02-03), was recovered from a nasopharyngeal swab taken from an RT-qPCR confirmed asymptomatic patient who returned from Wuhan, China in the beginning of February 2020. A close relation with the prototypic Wuhan-Hu-1 2019-nCoV (GenBank accession 112 number MN908947.3) strain was confirmed by phylogenetic analysis. Infectious virus was isolated by serial passaging on HuH7 and Vero-E6 cells (*36*); passage 6 virus was used for the study described here. The titer of the virus stock was determined by end-point dilution on Vero-E6 cells by the Reed and Muench method (*80*). The variant strain B.1.351 (hCoV-19/Belgium/rega-1920/2021; EPI_ISL_896474, 2021-01-11) was isolated from nasopharyngeal swabs taken from a traveler returning to Belgium and developing respiratory symptoms. The patients’ nasopharyngeal swabs were directly subjected to sequencing on a MinION platform (Oxford Nanopore) (Abdelnabi et al https://www.biorxiv.org/content/10.1101/2021.02.26.433062v1). Live virus-related work was conducted in the high-containment A3 and BSL3+ facilities of the KU Leuven Rega Institute (3CAPS), under licenses AMV 30112018 SBB 219 2018 0892 and AMV 23102017 SBB 219 20170589 according to institutional guidelines.

#### SARS-CoV-2 infection model in hamsters

Wildtype Syrian hamsters (Mesocricetus auratus) were purchased from Janvier Laboratories and were housed per two in ventilated isolator cages (IsoCage N Biocontainment System, Tecniplast) with ad libitum access to food and water and cage enrichment (wood block). Housing conditions and experimental procedures were approved by the ethical committee of animal experimentation of KU Leuven (license P065-2020). Female hamsters of 6-10 weeks old were anesthetized with ketamine/xylazine/atropine and inoculated intranasally with 50 μl containing 2×10^6^ or 1×10^4^ TCID50 for wt or B.1.351 variant, respectively. Treatment with mAb (human or hamster S2P6 (2-20 mg/kg) was initiated either 24) or 48 hours before infection by intraperitoneal injection. Hamsters were monitored for appearance, behavior and body weight. At day 4 post-infection, hamsters were euthanized by intraperitoneal injection of 500 μl Dolethal (200 mg/ml sodium pentobarbital, Vétoquinol SA). Lungs were collected, and viral RNA and infectious virus were quantified by RT-qPCR and end-point virus titration, respectively. Blood samples were collected before infection for pharmacokinetics analysis.

#### SARS-CoV-2 RT-qPCR

Hamster tissues were collected after sacrifice and were homogenized using bead disruption (Precellys) in 350 μl RLT buffer (RNeasy Mini kit, Qiagen) and centrifuged (10,000 rpm, 5 minutes) to pellet the cell debris. RNA was extracted according to the manufacturer’s instructions. To extract RNA from serum, a NucleoSpin kit (Macherey-Nagel) was used. 4 μl out of 50 μl eluate were used as a template in RT-qPCR reactions. RT-qPCR was performed on a LightCycler96 platform (Roche) using the iTaq Universal Probes One-Step RTqPCR kit (BioRad) with N2 primers and probes targeting the nucleocapsid (*36*). Standards of SARS-CoV-2 cDNA (IDT) were used to express viral genome copies per mg tissue or per ml serum.

#### End-point virus titrations

Lung tissues were homogenized using bead disruption (Precellys) in 350 μl minimal essential medium and centrifuged (10,000 rpm, 5 minutes, 4°C) to pellet the cell debris. To quantify infectious SARS-CoV-2 particles, endpoint titrations were performed on confluent Vero-E6 cells in 96-well plates. Viral titers were calculated by the Reed and Muench method (*80*) using the Lindenbach calculator and were expressed as 50% tissue culture infectious dose (TCID50) per mg tissue.

## Supplementary Information

**Table S1:** CryoEM data collection and refinement statistics.

|  | S2P6/SARS-CoV-2 S<br>(C3 map, post polishing) | S2P6/SARS-CoV-2 S<br>(C1 map, before polishing) |
| --- | --- | --- |
| **Data collection** |  |  |
| Magnification |  | 105,000 |
| Voltage (kV) |  | 300 |
| Total exposure (e<sup>-</sup>/Å<sup>2</sup>) |  | 60 |
| Defocus range (μm) |  | -0.1 to -5 |
| Pixel size (Å) |  | 1.201 |
| Initial particle stack |  | 1,092,473 |
| Final particle stack |  | 81,519 |
| Map resolution (0.143 FSC threshold) (Å) |  | 4.2 |
| Map B-factor |  | 102.2 |
| Symmetry |  | C1 |

**Table S2:**
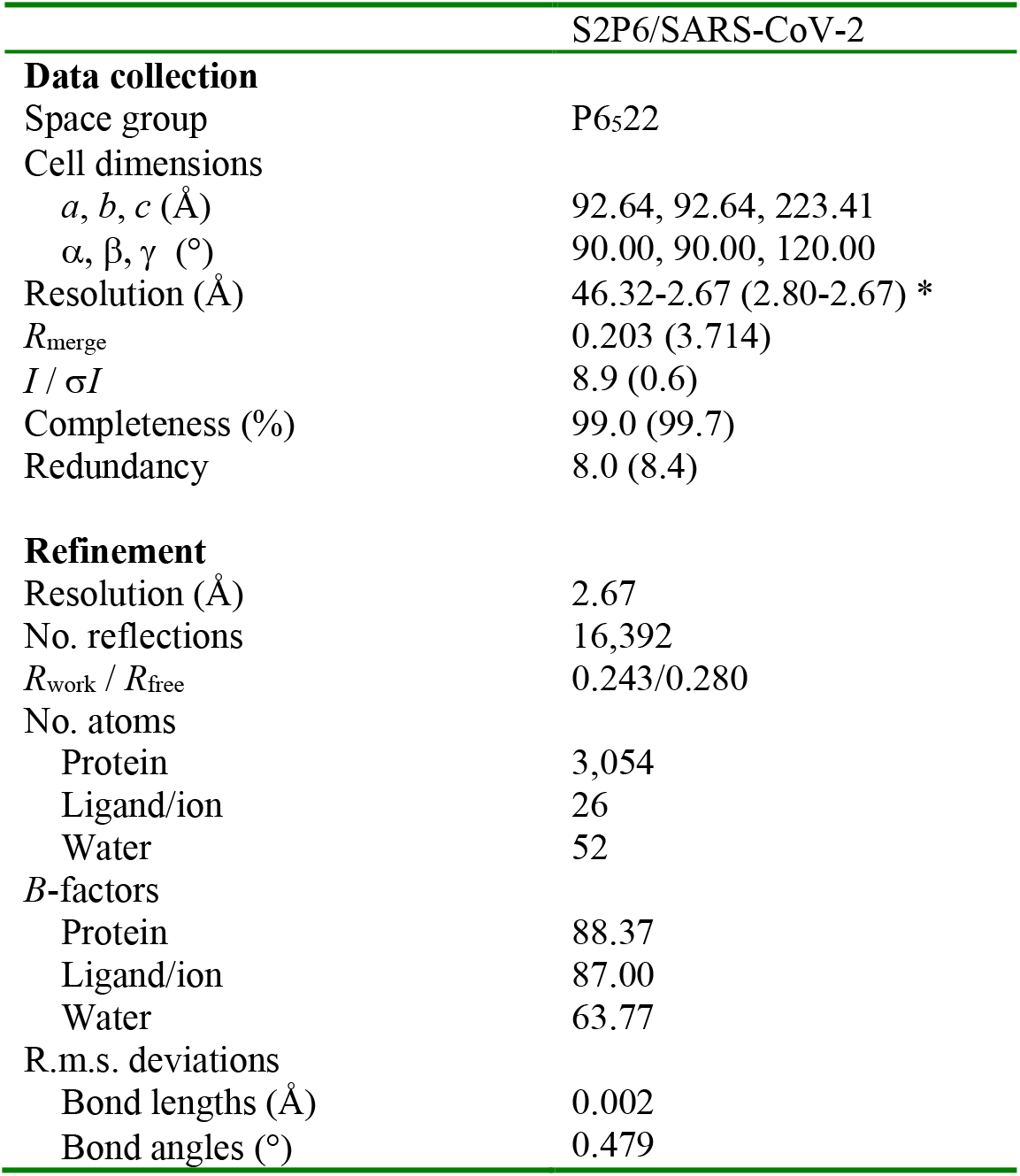
X-ray crystallography data collection and refinement statistics.

|  | S2P6/SARS-CoV-2 |
| --- | --- |
| **Data collection** |  |
| Space group | P6<sub>5</sub>22 |
| Cell dimensions |  |
| *a*, *b*, *c* (Å) | 92.64, 92.64, 223.41 |
| α, β, γ (°) | 90.00, 90.00, 120.00 |
| Resolution (Å) | 46.32-2.67 (2.80-2.67) * |
| *R*<sub>merge</sub> | 0.203 (3.714) |
| *I* / σ*I* | 8.9 (0.6) |
| Completeness (%) | 99.0 (99.7) |
| Redundancy | 8.0 (8.4) |
| **Refinement** |  |
| Resolution (Å) | 2.67 |
| No. reflections | 16,392 |
| *R*<sub>work</sub> / *R*<sub>free</sub> | 0.243/0.280 |
| No. atoms |  |
| Protein | 3,054 |
| Ligand/ion | 26 |
| Water | 52 |
| *B*-factors |  |
| Protein | 88.37 |
| Ligand/ion | 87.00 |
| Water | 63.77 |
| R.m.s. deviations |  |
| Bond lengths (Å) | 0.002 |
| Bond angles (°) | 0.479 |

**Fig. S1.**
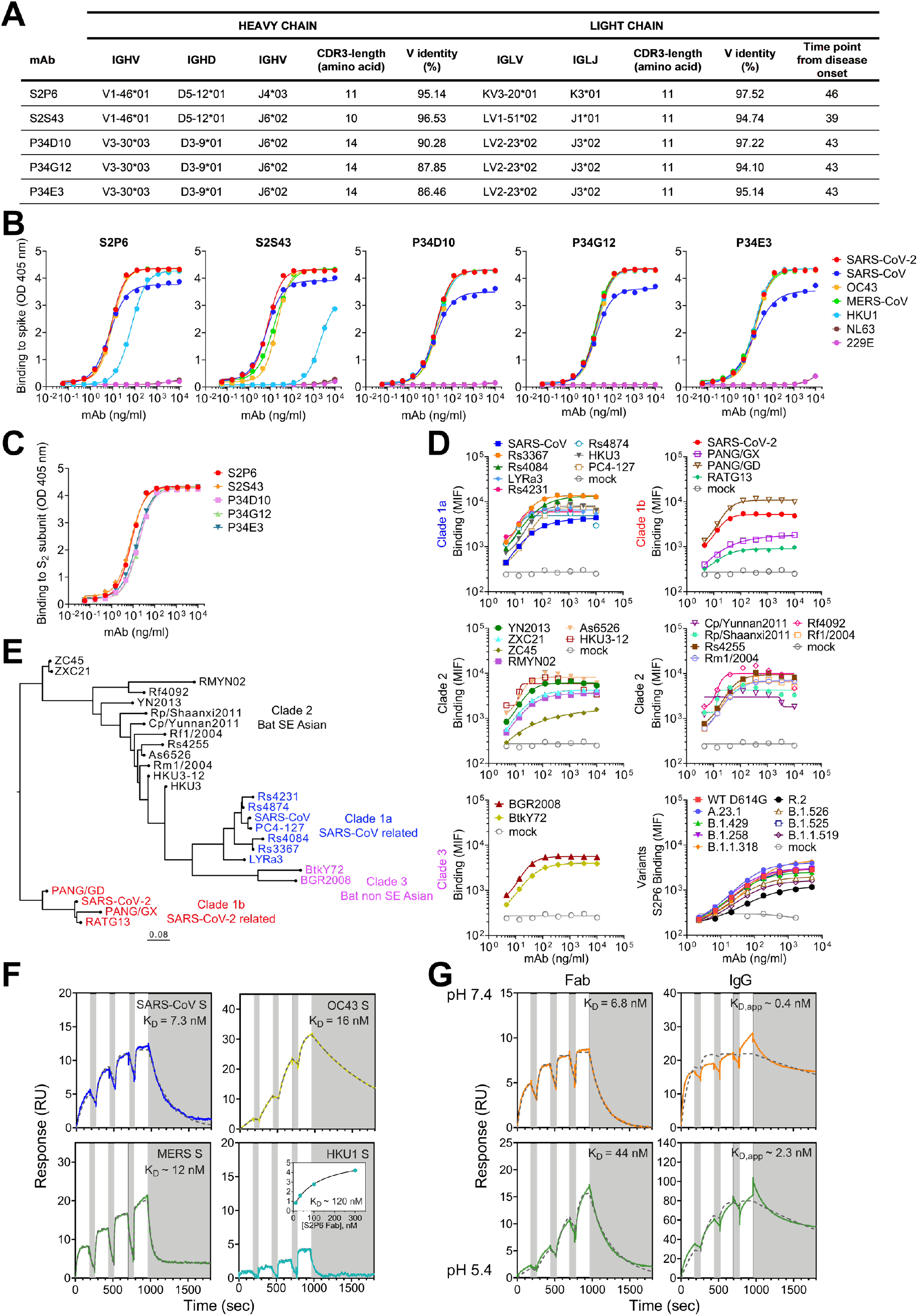
Properties of the 5 cross-reactive mAbs isolated. (**A**) V(D)J usage, nucleotide sequence identity to germline genes, number of somatic mutations, and time interval between sample collection and mAb isolation. (**B**) Binding of identified mAbs to prefusion β-coronavirus S ectodomain trimers by ELISA (**C**) Binding to SARS-CoV-2 post-fusion S_2_ subunit. (**D**) Mean fluorescence intensity as measured in flow cytometry for S2P6 binding to a panel of 26 S glycoproteins representative of all *sarbecovirus* clades and 8 SARS-CoV-2 variants. (**E**) Phylogenetic tree of sarbecoviruses S used in this work inferred via maximum likelihood analysis of spike amino acid sequences. (**F**) SPR analysis of S2P6 Fab binding to immobilized prefusion β-coronavirus S trimers. (**G**) SPR analysis of S2P6 Fab and IgG binding to immobilized prefusion SARS-CoV-2 S ectodomain trimer at pH 7.4 and pH 5.4. Fits to a 1:1 binding model are an approximation for IgG binding due to bivalency.

**Fig. S2.**
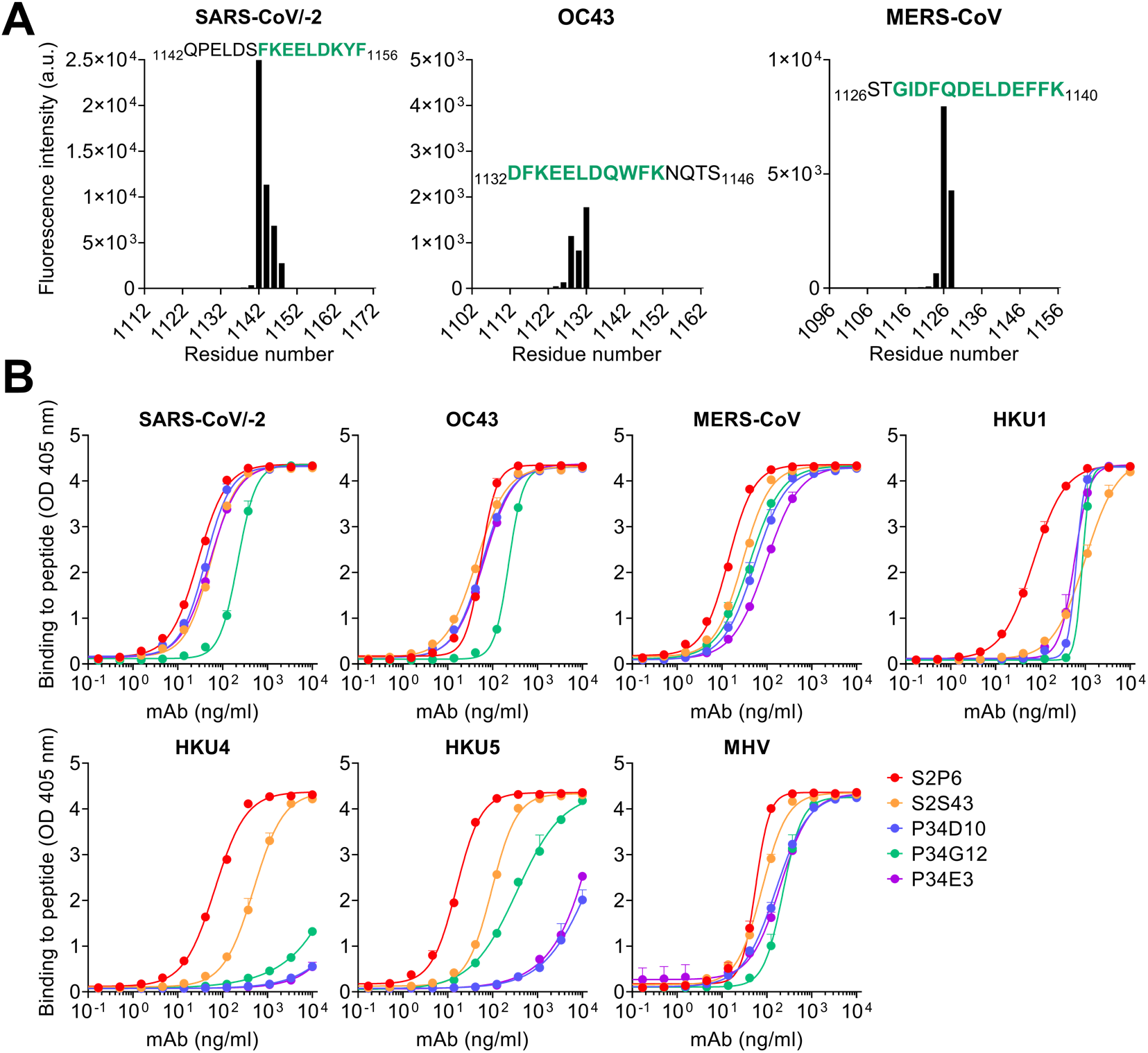
Identification of the S2P6 epitope. (**A**) Binding of S2P6 to linear peptides (15-mer peptides overlapping by 13 residues) spanning the SARS-CoV/SARS-CoV-2 S, OC43 S and MERS-CoV S sequences. (**B**) Binding of identified mAbs to β-coronavirus S stem helix peptides by ELISA.

**Fig. S3.**
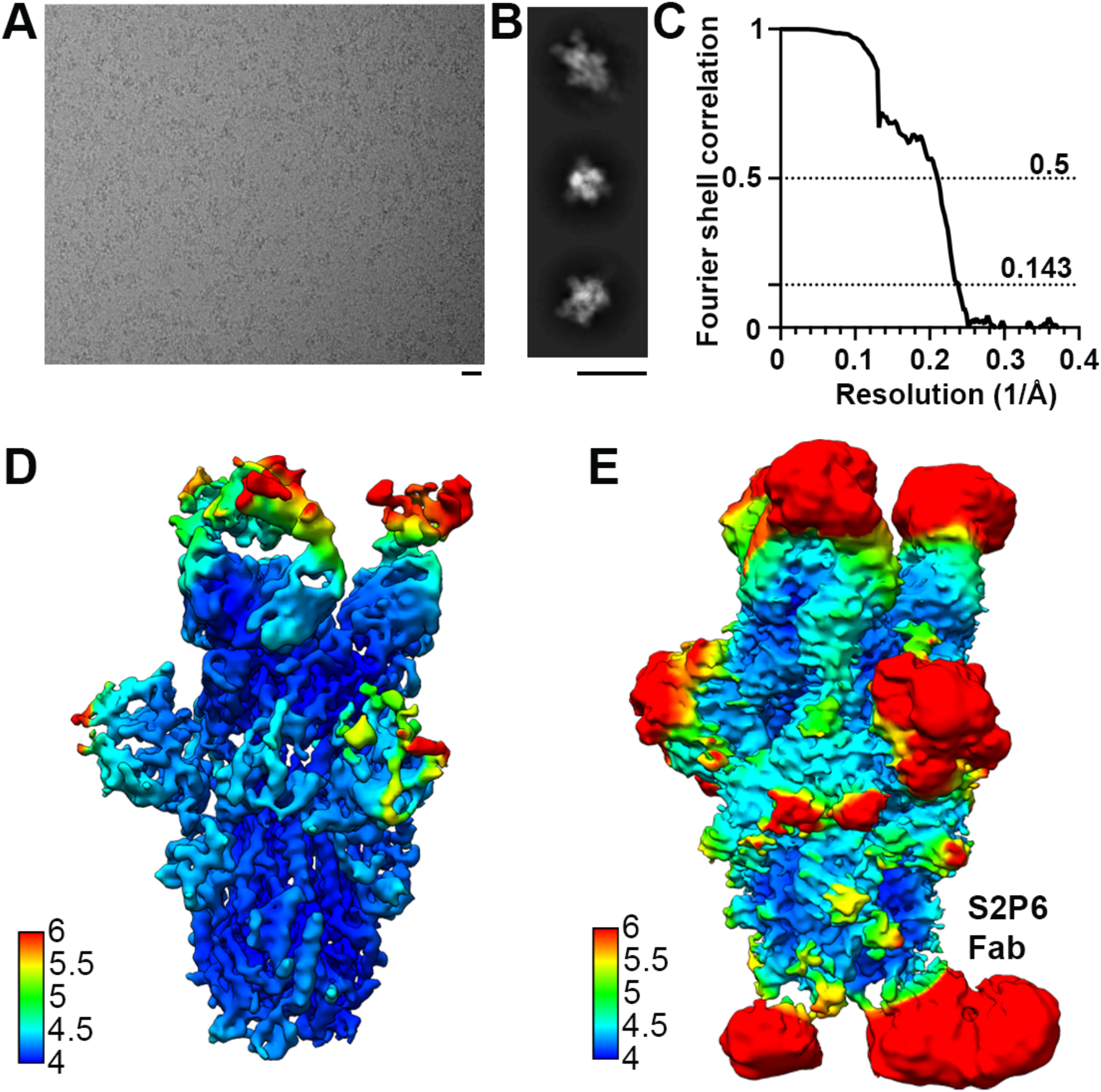
CryoEM data processing and validation of S2P6- and S2M11-bound SARS-CoV-2 S dataset. (**A-B**) Representative electron micrographs (A) and class averages (B) of SARS-CoV-2 S in complex with the S2P6 and S2M11 Fabs. Scale bars: 200 Å. (**C**) Gold-standard Fourier shell correlation curve. The 0.143 and 0.5 cut-offs are indicated by horizontal dashed gray lines. (**D-E**) CryoEM map colored by local resolution computed using cryoSPARC shown at two distinct contour levels.

**Fig. S4.**
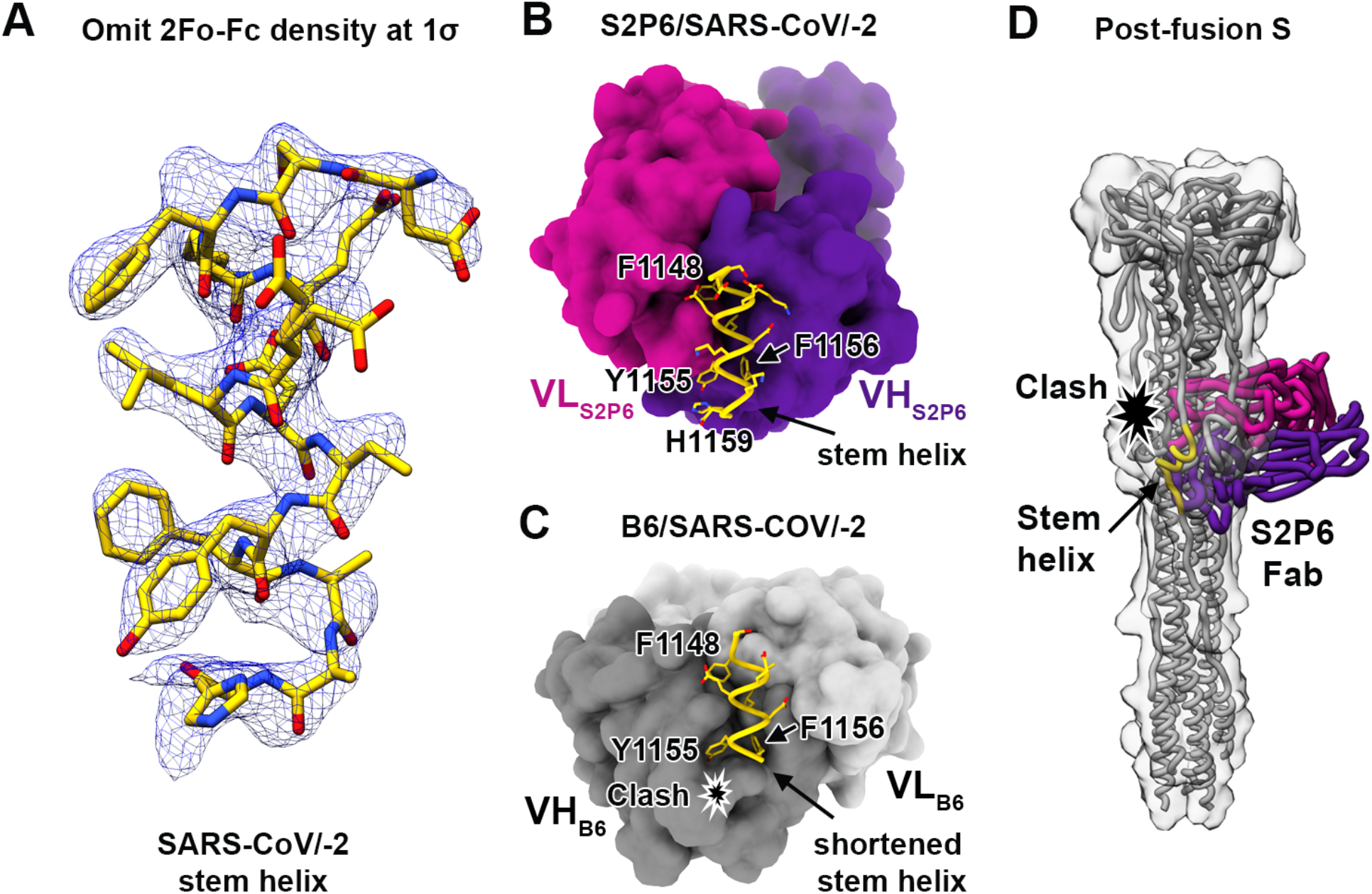
Comparison of the S2P6 and B6 mAb binding modes. (**A**) Crystal structure of the SARS-CoV-2 stem helix peptide rendered as sticks with the corresponding 2Fo-Fc omit map contoured at 1.5σ. The S2P6 Fab fragment is not shown for clarity. (**B**) Crystal structure of the S2P6 Fab (surface rendering) in complex with the SARS-CoV-2 S stem helix peptide (yellow ribbon with side chains rendered as sticks). (**C**) Crystal structure of the B6 Fab (surface rendering) in complex with the SARS-CoV-2 S stem helix peptide (yellow ribbon with side chains rendered as sticks). The star indicates the putative clash between B6 CDRH2 and the stem helix C-terminus, likely explaining the latter region is disordered in the B6-bound structure whereas it is resolved in the S2P6-bound structure. (**D**) Superimposition of the S2P6-bound (purple/magenta) SARS-CoV-2 stem helix (yellow) crystal structure onto the SARS-CoV S post-fusion structure (PDB 6M3W) shows that S2P6 binding would be incompatible due to steric hindrance suggesting S2P6 prevents S fusogenic conformational changes. A low-pass filtered surface generated from the SARS-CoV S post-fusion structure is shown as a transparent gray surface to help visualizing clashes.

**Fig. S5.**
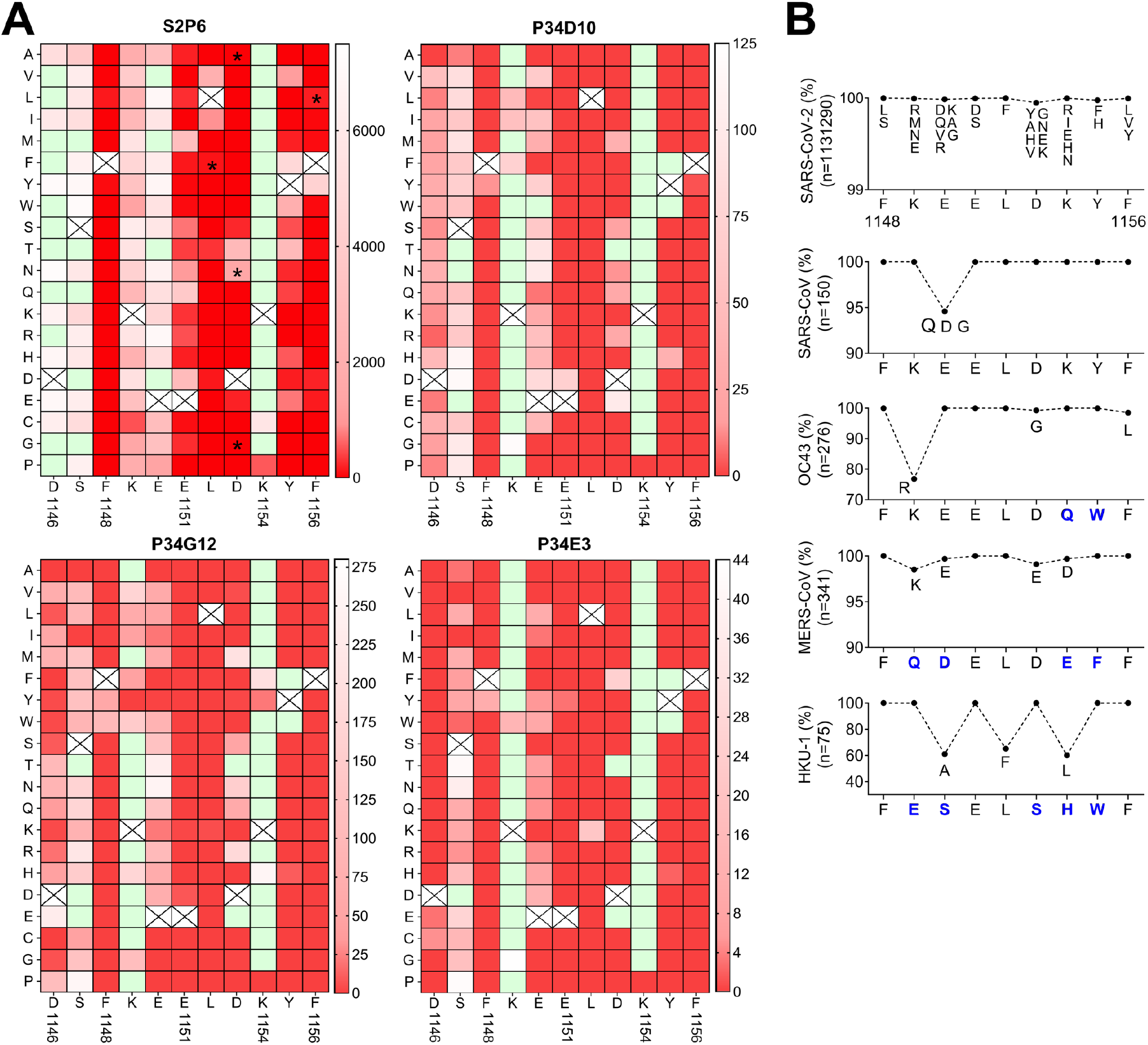
Impact of individual SARS-CoV-2 stem helix residue substitution on mAb binding. (**A**) Heat map showing binding (fluorescence intensity) of S2P6, P34D10, P34G12 and P34E3 to stem helix peptides harboring each possible amino acid substitution. White to red gradient indicate the degree of loss of binding as compared to the native residue (white) shown as a crossed square. Green squares indicate substitutions enhancing binding as compared to the native residue. Asterisks highlight viral escape substitutions identified in vitro for S2P6. (**B**) Epitope conservation among β-coronavirus sequences with human and animal hosts retrieved from GISAID. The consensus sequence for SARS-CoV-2 is reported on x axis and predominant substitutions are indicated by a blue letter.

**Fig. S6.**
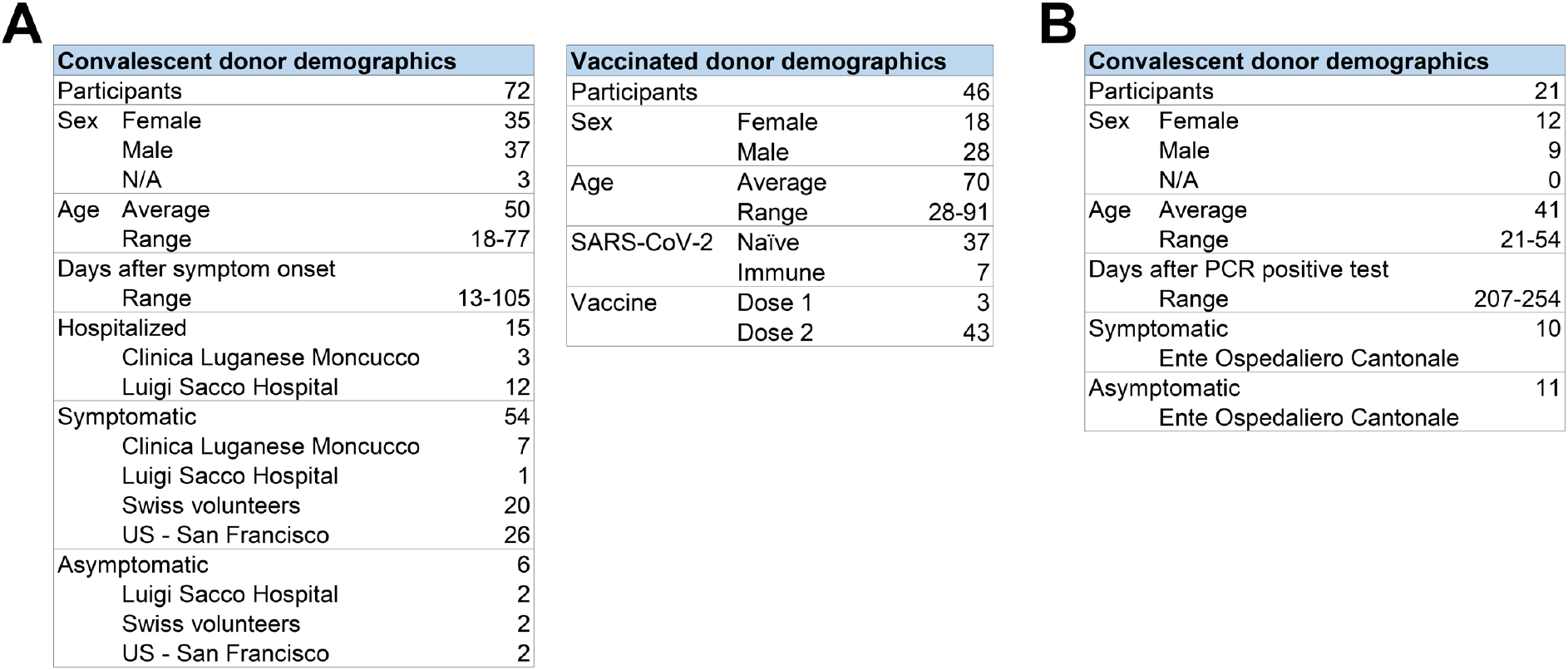
Patient demographics. (**A**) Summary of convalescent patient demographics from which plasma (left table) or memory repertoire (right table) have been analyzed. (**B**) Summary of vaccinees patient demographics.

**Fig. S7.**
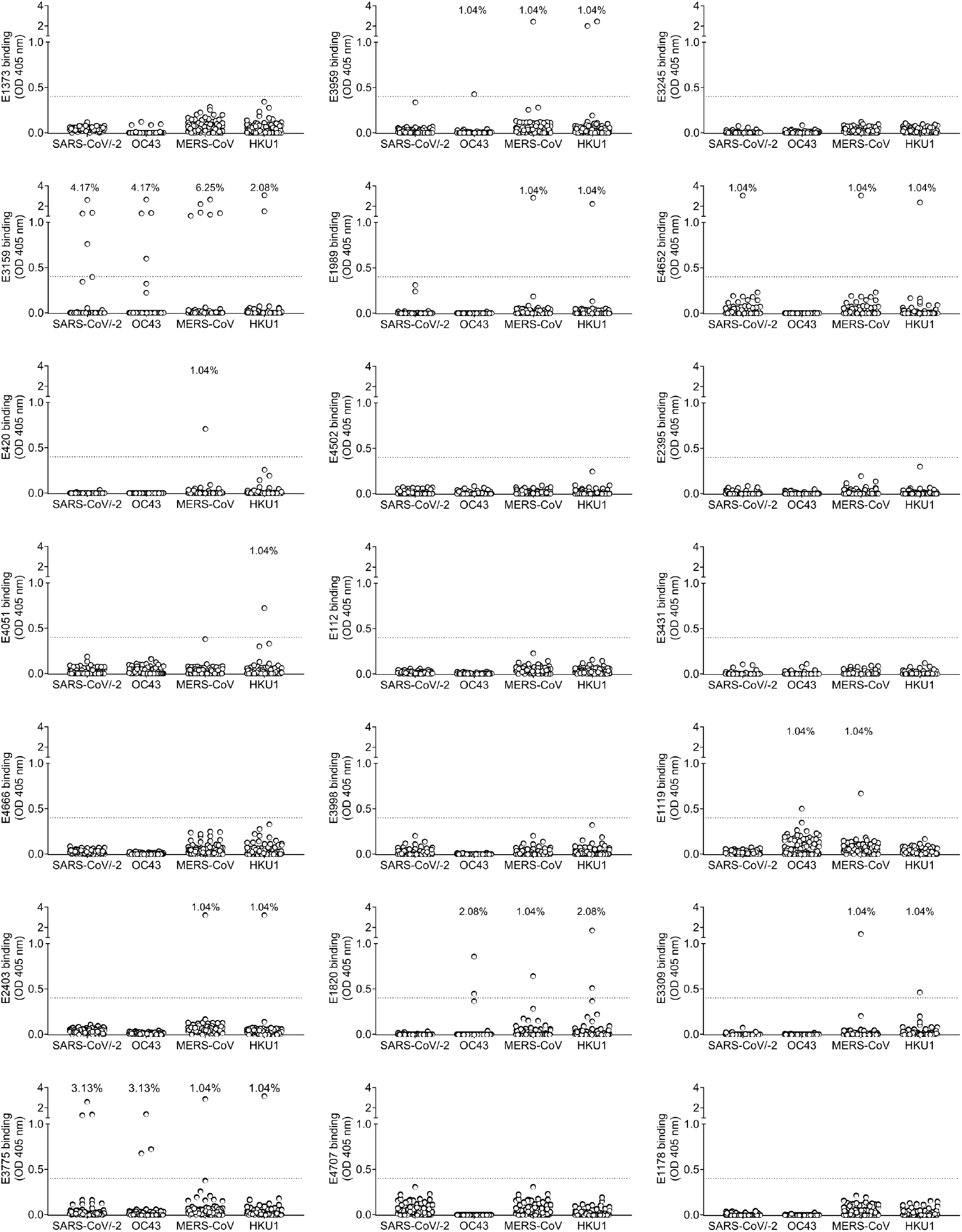
Binding of IgG memory B-cells from COVID-19 convalescent individuals to β-coronavirus stem helix peptides. Cut-off (OD=0.4) is indicated by a dotted line and frequencies of cultures scoring positive are reported for each antigen.

**Fig. S8.**
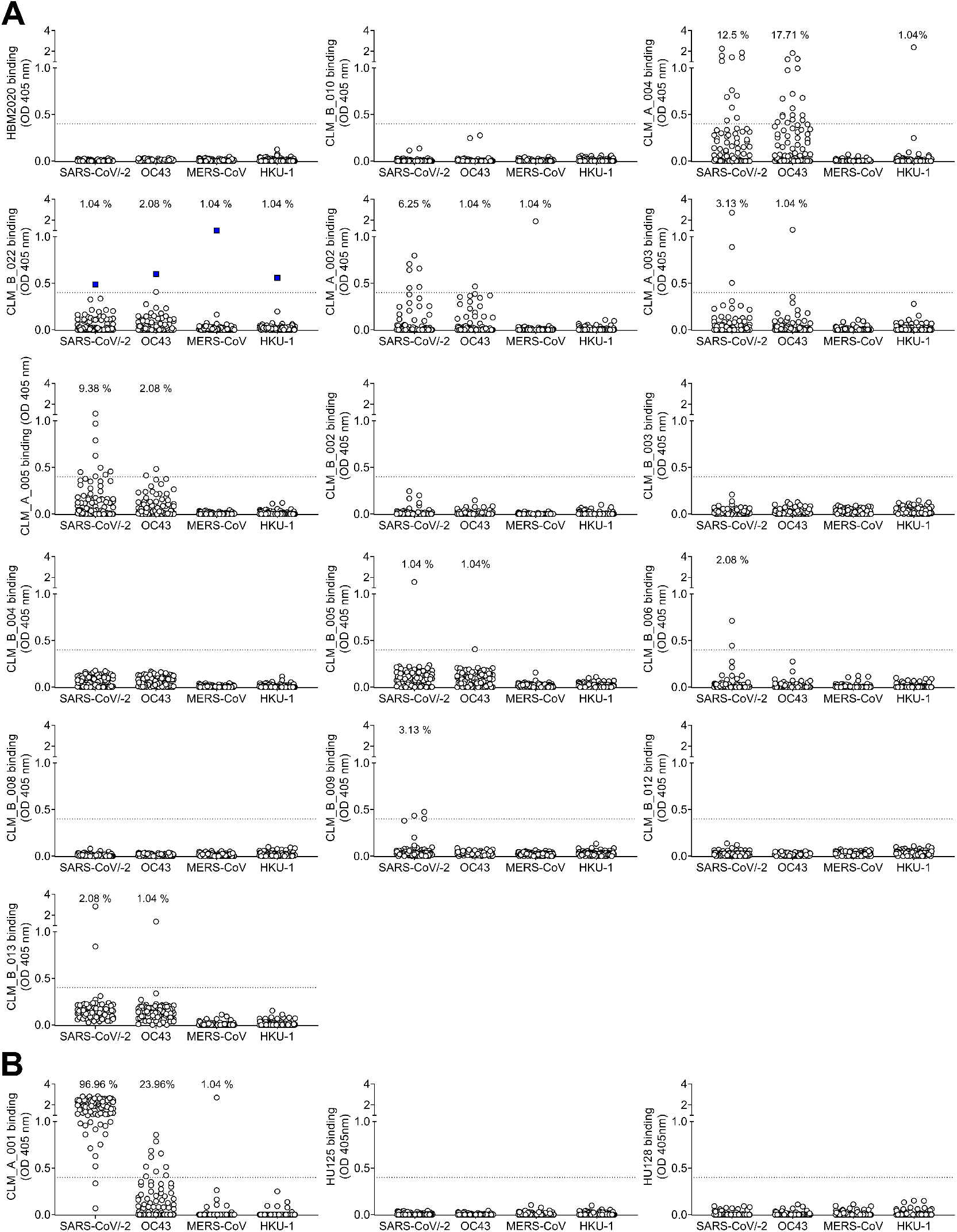
Binding of IgG memory B-cells from COVID-19 vaccinees to β-coronavirus stem helix peptides. (**A**) Cut-off (OD=0.4) is indicated by a dotted line and frequencies of cultures scoring positive are reported for each antigen. (**B**) Binding to stem helix peptide of a previously infected individual after first vaccine dose showing high response to SARS-CoV-2 (left) and of two pre-pandemic individuals (middle and right).

**Fig. S9.**
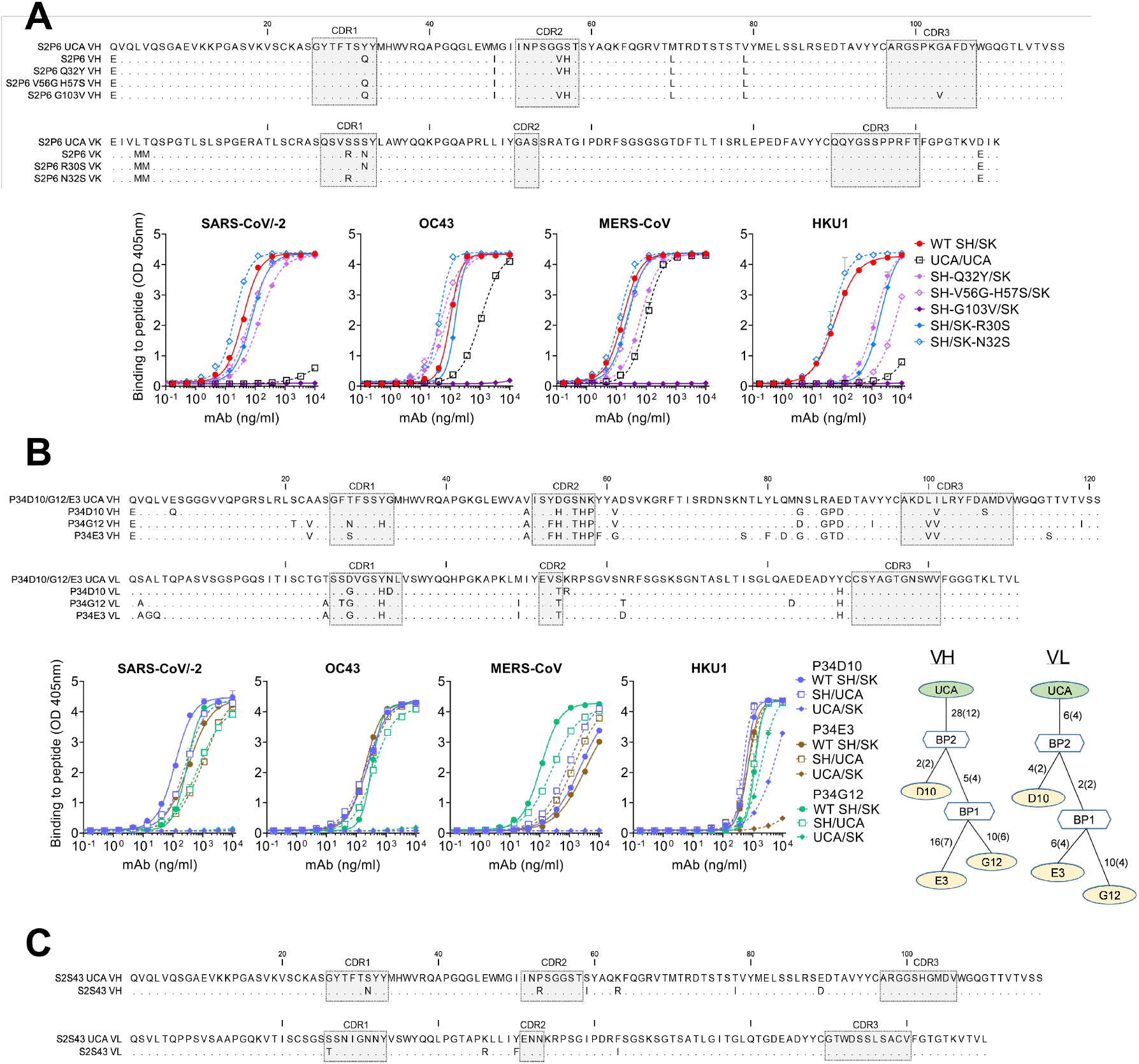
Analysis of mAbs sequence. (**A**) Alignment of the amino acid sequences of S2P6 UCA and wild-type VH and VL (top). The CDRs regions are highlighted in gray. Binding of S2P6 mutated SH or SK with fully germline reverted residues in CDR (bottom). (**B**) Alignment of the amino acid sequences of P34D10, P34G12 and P34E3 UCA and wild-type VH and VL (top). Binding of mAbs mature heavy chain paired with germline reverted light chain (SH/UCA) and germline reverted heavy chain paired with mature light chain (UCA/SK) (bottom). Genealogic trees of VH and VL genes of P34D10, P34G12 and P34E3 shown using AncesTree (*81*). The number of nucleotide and amino acid (in parentheses) mutations from the unmutated common ancestor (UCA) or branch points (BPs) to their descendants are shown. (**C**) Alignment of the amino acid sequences of S2S43 UCA and wild-type VH and VL

